# Phage N4 uses a SAR endolysin-holin system for host cell lysis

**DOI:** 10.1101/2025.11.12.688109

**Authors:** Michael B. Awuah, Cody Martin, Jake S. Chamblee, Adam J. Tomaszewski, Teresa E. Sullivan, Qori Emilia, Steven Tran, Jason H. Snowden, Kaylyn A. Niemiec, Jiatong Zhu, Jolene Ramsey

## Abstract

Bacteriophages (phages) cause active host cell lysis to terminate their infection and release progeny into the environment. However, some phages have systems to delay their lysis, and thereby increase progeny yield, in specific conditions: a phenomenon called lysis inhibition (LIN). Two dissimilar phages of *Escherichia coli* are known to exhibit LIN: T4 and N4. In T4, a multi-protein mechanism stalls lysis and maintains the LIN state in response to superinfection in a high phage population density. However, the lysis proteins responsible for T4 lysis and LIN are not conserved with phage N4. In this study, we characterize the phage N4 proteins involved in lysis by molecular and genetic means. We define the functions of the minimal gene set required for lysis through heterologous expression and complementation. Furthermore, by sequence comparison with a selected mutant library that does not induce LIN, we have identified genomic regions both within and outside the lysis cassette involved in N4 LIN. We propose a model where N4 lysis proteins that execute rapid N4 lysis can be regulated to induce LIN. Despite the lack of conservation with T4 components, our study suggests that direct modulation of lysis initiation may be common and provides a springboard for identifying similar mechanisms in other phages.

**Importance:** Phages are viruses that kill only bacteria. To do so, they use lysis proteins to disrupt the structural components of the bacterial envelope. There are commonalities between the phages that infect bacteria with similar structures, but there are also interesting differences we can exploit to learn more about specific phages or their bacterial hosts. In this study, we characterize the lysis proteins of the *E. coli* phage N4. Our results suggest that lysis regulation controls phage yield in N4 differently than in well-studied phage T4. Future studies on the relationship between lysis proteins and phage yield could provide handles to use this information for optimal large-scale phage production for clinical or industrial applications.

## Introduction

Bacteriophages (phages) are viruses that infect and kill bacteria via a process known as lysis. Characterization of phage-mediated host cell lysis in the golden age of phage biology revealed a multi-gene lysis (MGL) paradigm common among tailed phages (1). Variations within their MGL systems at the mechanistic and regulatory levels afford phages evolutionary flexibility (2–5). For example, coliphage N4, whose large, packaged virion RNA polymerase (vRNAP) is extensively and elegantly described from a biochemical perspective, increases its progeny yield an order of magnitude higher via an unknown lysis regulation pathway (6–8). As a tailed phage with a 72-kb dsDNA genome, phage N4 is expected to encode a full suite of MGL proteins for progeny release.

A minimal set of MGL proteins includes the holin for initiating lysis and the endolysin for cell wall degradation (9). In bacterial cells with additional cell envelope layers, such as the mycomembrane of mycolata hosts or the outer membrane of diderms, specialized proteins like the phage spanins found in phages that infect Gram-negative bacteria are necessary for lysis to allow progeny release (10–13). There are two major MGL systems in phages of Gram-negative bacteria: the holin-endolysin-spanin system and the pinholin-SAR endolysin-spanin system (1). In the canonical phage λ holin-endolysin-spanin system, pore-forming proteins known as holins accumulate harmlessly within the inner membrane. Upon reaching an allele-specific critical threshold concentration, holins depolarize the inner membrane by making micron-sized holes (14, 15). Hole formation gives the muralytic endolysins access to their cell wall substrate in the periplasmic space. Degradation of the peptidoglycan meshwork liberates fusogenic proteins known as spanins to fuse the inner and outer membrane, leading to cell envelope rupture (16). Similarly, in the phage 21 pinholin-SAR endolysin-spanin system, nanometer-scale pinholes depolarize the inner membrane (17). Pinholin action simultaneously releases signal-arrest-release (SAR) endolysins into the periplasm where they assume an active conformation (18). Due to the weakly hydrophobic nature of the SAR domain, SAR endolysins also stochastically leak from the inner membrane. When sufficient peptidoglycan is degraded, spanin action leads to explosive cell lysis. Most characterized tailed dsDNA phages infecting Gram-negative hosts encode an MGL system with these functions to release progeny across the cell envelope.

The classic phage T4, a representative of the T-even phages, encodes a canonical MGL system and dynamically increases its progeny yield by regulating the holin (19). The T4 holin, called T, has a transmembrane domain (TMD) that anchors a large periplasmic domain (20). When the T periplasmic domain is bound by the regulatory antiholin protein rI, hole formation is delayed (21, 22). By delaying hole formation, lysis time is delayed for hours, a phenomenon known as lysis inhibition (LIN) (19). In T4, a series of LIN-defective rapid lysis mutants (*r*) were identified based on a change in their plaque morphology (23). Those early genetic studies found that mutations in the T4 holin gene (*rV*), two antiholins (*rI* and *rIII*), and a periplasmic protein called spackle (*gp61.3*) displayed LIN *r* plaques (24). Later biochemical and structural studies demonstrated that the LIN system of phage T4 hinges on the direct holin protein inhibition by antiholins (21, 22, 25). The holin and LIN genes are conserved within closely related T-even phages of *E. coli* (19). Holin regulation has also been observed in T4-like phages that infect *Vibrio cholerae* where there is no sequence similarity (26). Therefore, holin regulation seems to represent a common lynchpin between lysis and LIN pathways.

Despite holin’s functional conservation across MGL systems, the holin proteins exhibit high sequence variation. In fact, without sequence similarity or genetic evidence, holins are typically discovered by experimental characterization of genes encoding a transmembrane domain. Therefore, detailed lysis studies are imperative to construct an accurate picture of the lysis landscape across phage and host cells. In phage N4, spanins were bioinformatically identified and an active endolysin was reported in 2007, but holins are only predicted (27, 28). Moreover, the pathway leading to the N4 LIN phenotype remains unstudied (29, 30). Wildtype (WT) N4 displays asynchronous lysis in bulk culture starting ∼3 hours after infection, its typical LIN phenotype (7). When LIN was observed in N4 in the late 1960’s, Schito and colleagues who isolated N4 *r*, a variant that completes lysis within an hour after infection (7, 30). However, the allele responsible for the N4 *r* phenotype was not mapped or sequenced, leaving the genetic basis of N4 LIN unidentified. Without identified holins, phage N4 lacks a molecular framework for LIN studies. Additionally, the past decade has witnessed a boom in reports of N4-like phages defined by encoding the characteristic vRNAP (31–33). Most N4-like phage reports do not mention LIN at all. Although N4-like *Pseudomonas* phage AM.P2 may exhibit LIN, *E. coli* phage G7C and *Achromobacter* phages JWDelta and JWAlpha do not exhibit LIN (34–36). To better understand lysis and LIN within the N4-like phages, the holin of the phage N4 lysis system must be identified.

Here, we demonstrate that phage N4 encodes a complete and functional MGL system, including two holin proteins. Heterologous expression of each N4 lysis protein provides experimental evidence of their individual contributions to lysis. Expression of the full N4 MGL system alone leads to rapid lysis. The N4 master regulator pinholin and its second holin protein usually co-occur in N4-like phages, suggesting commonalities in their lysis regulation pathways.

## Results

### The phage N4 predicted lysis genes cause rapid host lysis

Seeking further evidence for a LIN system in N4, we imaged N4-infected cells at times post-infection when population lysis was ongoing (Fig. 1A). As previously reported using electron microscopy (29), many cells were elongated and roughly rod shaped (Fig. S1), suggesting that the endolysin had not destroyed the PG. At 30 min post-infection, N4 WT-infected cells were already longer at 7.6 ± 2.3 µm compared to uninfected cells at 4.5 ± 1.1 µm. Cell enlargement in infected cultures continued progressively through 120 min post-infection, reaching mean values of 12.5 ± 3.6 µm in length and 2.4 ± 0.5 µm in width. In contrast, uninfected cells decreased in size over the same period, consistent with transitioning from exponential growth to stationary phase. Additionally, when individual cells lysed, they did so rapidly and completely (Fig. 1A, Movies S1 & S2). Before lysis, each rod-shaped cell contracted from the poles, characteristic of SAR endolysin activity degrading the peptidoglycan evenly around the cell body after release from the inner membrane (1). This lysis pattern suggests that individual N4-infected cells lyse with the standard pattern expected for complete MGL systems, what is referred to as rapid lysis.

**Figure 1.**
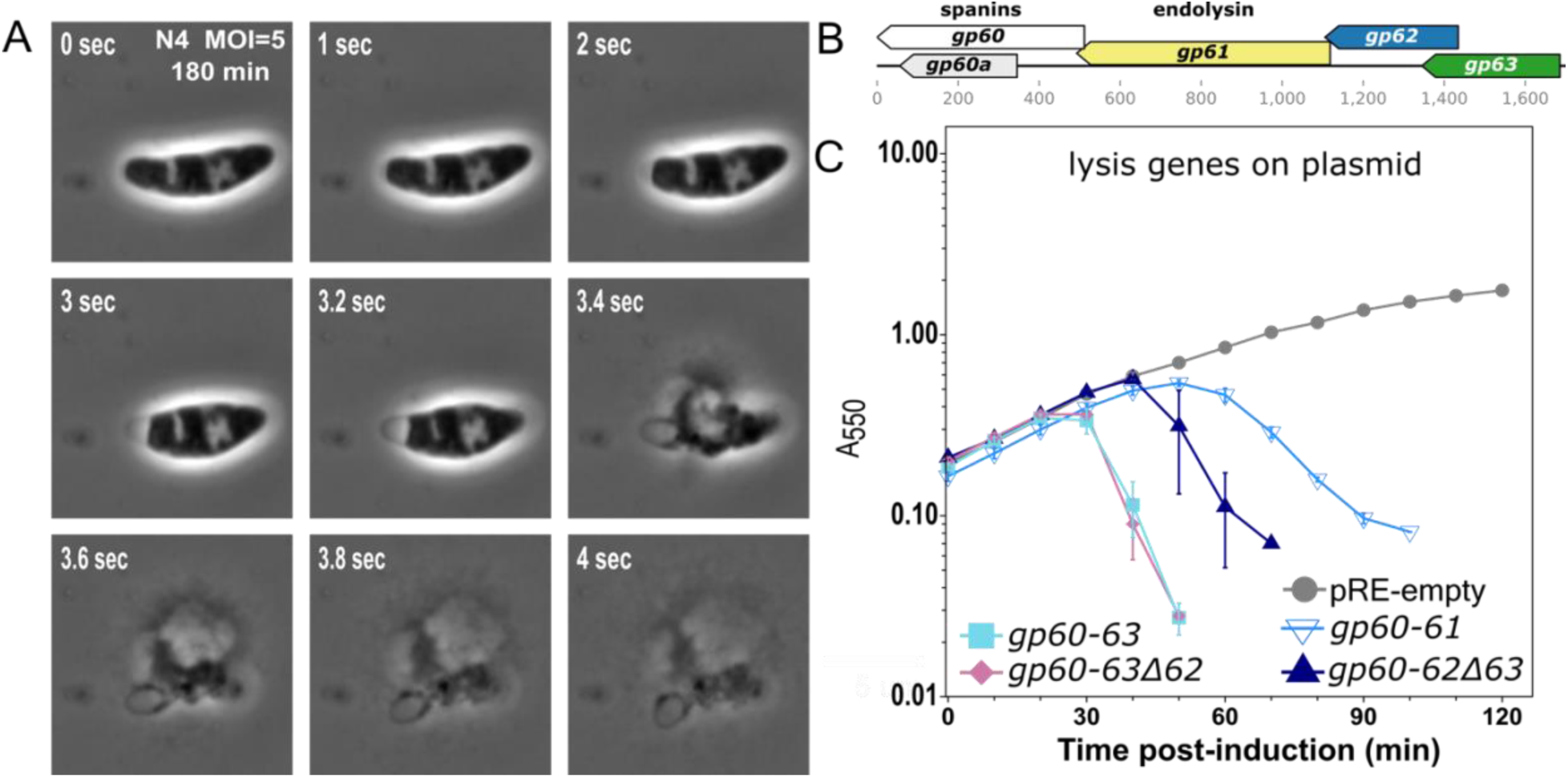
Phage N4 encodes a multi-gene lysis system capable of effecting rapid lysis. **A)** Representative time lapse microscopy of MG1655 180 minutes after infection with N4 WT at an MOI=5. The complete time lapse video is Movie S1. **B)** Genetic organization of the phage N4 lysis genes including the i-spanin *gp60*, the embedded o-spanin *gp60a*, the endolysin *gp61*, the holin *gp62*, and the pinholin *gp63*. **C)** Lysis kinetics of MG1655 cultures expressing N4 lysis genes from the pRE plasmid under the λ Q-dependent pR’ promoter. Host cultures harboring derivatives of pRE carrying N4 lysis genes and the pQ-Kan plasmid (λ Q under lacPO1 control) were induced with 1 mM IPTG at T=0 min. Lysis was monitored by optical density for pRE-empty (solid gray circle), *gp60-62Δ63* (solid navy blue triangle), *gp60-63* (solid cyan square), *gp60-63Δ62* (solid pink diamond), and *gp60-61* which includes embedded *gp60a* (open inverted blue triangle). Lysis curves represent at least three biological replicates and data are presented as the mean ± standard error of the mean.

Re-inspection of the phage N4 genome annotation alongside N4-like phages revealed clustered late genes encoding proteins with the classic characteristics of spanins and holins flanking the endolysin (Fig. 1B) (10). To demonstrate that the N4 WT genome encoded the genes necessary and sufficient to effect rapid lysis, we cloned the putative lysis cassette *gp60*-*gp60a*-*gp61*-*gp62*-*gp63* (abbreviated *gp60-63*) into an inducible plasmid expression vector under the control of the lambda late promoter, which achieves expression levels known to be effective for coordinated rapid lysis in *E. coli* (37–39). Upon induction, we observed culture lysis at ∼25 minutes (Fig. 1C), indicating that the *gp60-63* gene cluster constitutes a lysis cassette encoding a complete MGL pathway. To evaluate contribution of each gene to lysis onset, defined as the time when optical density (OD) begins to drop, and lysis completion, defined as the time from lysis onset to an OD drop below 0.1, we constructed a series of gene deletions. The *gp62* deletion did not change the lysis pattern, suggesting that gp62 plays a nonessential role in lysis. The *gp63* deletion resulted in a 20-minute delay to lysis onset and completion. The double *gp62* and *gp63* deletion (known as *gp60-61*) led to gradual culture lysis from 50-90 minutes post-induction (Fig. 1C). These data suggest that gp63 is the master regulator of lysis onset. Additionally, these data suggest that gp62 is sufficient to induce sudden lysis when gp63 is absent. To further delineate the molecular roles of each gene in the lysis cassette, we proceeded to characterize their molecular properties.

### N4 encodes a SAR endolysin and bimolecular spanins

The cellular morphology shift during lysis and slow completion of lysis in the presence of gp60-61 (Fig. 1C) is reminiscent of a SAR endolysin phenotype. Indeed, Stojković and Rothman-Denes demonstrated that purified gp61 is a membrane-associated N-acetylmuramidase in 2007 (27). Inspection of the gp61 N-terminal sequence revealed a weakly hydrophobic domain rich in glycine and alanine and flanked by charged lysine residues, the characteristics of a SAR domain (shown with Alphafold3 prediction in Fig. S2). As expected, gp61 complemented cell wall disruption activity when expressed in lysogens encoding either a canonical large hole or a pinholin but no other endolysin (Fig. 2A-B). Functioning with the pinholin suggested that gp61 was a SAR endolysin. Gp61 expressed alone demonstrates low levels of cell growth inhibition, comparable to the levels observed with the phage 21 SAR endolysin, on a timescale that matches asynchronous exit from LIN (Fig. 2C). Microscopy revealed that the morphology of rod-shaped cells observed at the start of the experiment were converted to round morphology for R^21^ and partially converted for gp61 (Fig. 2C). The gradual morphology change is consistent with known asynchronous SAR endolysin stochastic release from the membrane, in addition to their typical holin-dependent synchronous release (18). We conclude that gp61 is a SAR endolysin.

**Figure 2.**
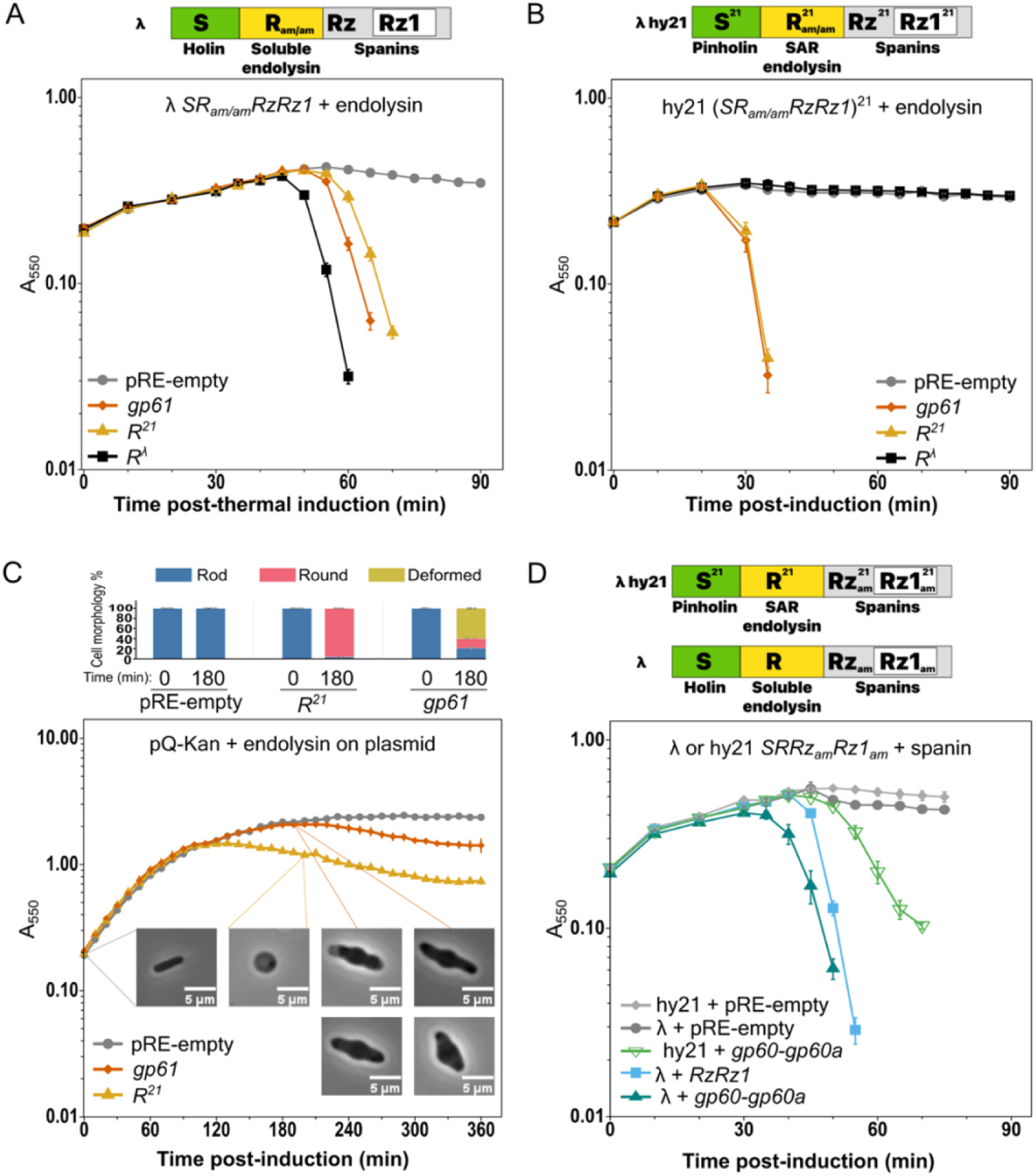
N4 endolysin and spanin function in lysis. Lysis gene activity in the **A)** λ lysogen with an amber allele in the soluble endolysin and **B)** hy21 lysogen with an amber allele in the SAR endolysin was monitored after thermal induction in the presence of pRE-empty (solid gray circle), N4 *gp61* (solid orange diamond), phage 21 endolysin *R^21^* (solid gold triangle), or λ endolysin *R^λ^* (solid black square). The hy21 lysogen is concurrently induced with IPTG. **C)** Lysis kinetics of MG1655 cultures with pRE -empty (solid gray circle), N4 *gp61* (solid orange diamond), or *R^21^*(solid gold triangle) plasmids. Rod-shaped (blue), round (pink), or deformed (gold) cell morphologies at T = 0 min and T = 180 min following induction were assessed through phase contrast microscopy. Representative micrographs for each morphological category are shown (scale bars, 5 µm). Open circles represent values from independent biological replicates (n = 3) with means calculated from pRE -empty n=856, 1191 cells, *gp61* n=542, 933 cells, and *R^21^* n=856, 379 total cells counted at 0 and 180 minutes, respectively for each. **D)** As in A and B, except the lysogens had amber alleles in the spanin genes. Lysis gene activity was monitored for pRE-empty (solid gray diamonds and circles), λ spanins (*RzRz1*) (solid blue square), or N4 spanin *gp60* and *gp60a* (open and solid green triangles). All lysis curves represent at least three biological replicates and data are presented as the mean ± standard error of the mean.

The genes downstream of *gp61* were identified bioinformatically as the type members of a large spanin gene family (10, 28). The *gp60* inner membrane spanin (i-spanin) gene is predicted to encode a type II single-pass integral membrane protein, with 140 residues in the periplasm (shown with an Alphafold3 prediction in Fig. S3). The *gp60a* outer membrane spanin (o-spanin) gene is predicted to encode a mature outer membrane lipoprotein of 56 residues (shown with an Alphafold3 prediction in Fig. S3). As is the case for many bimolecular spanins, the *gp60a* open reading frame is fully embedded within the *gp60* gene. Upon expression during induction of a spanin-defective λ lysogen, N4 gp60-gp60a exhibited full complementation of the outer membrane disruption defect (Fig. 2D), indicating they are functional bimolecular spanins.

### N4 encodes a pinholin and holin

Based on the canonical MGL model, N4 is anticipated to initiate lysis via a master regulator holin. Other than having nondescript TMDs, holins lack conserved domains, necessitating experimental validation to definitively assign function. The two TMD-containing proteins encoded upstream of the SAR endolysin are therefore holin candidates. The *gp63* gene encodes a protein of 110 residues with an alternate start site at Met17. In contrast to most type II integral membrane proteins, where the TMD is located at the N-terminus, full-length gp63 is predicted to be C-terminally anchored in the inner membrane (shown with an Alphafold3 prediction in Fig. S4). Although various TMD prediction tools report both possible orientations, following the positive-inside rule we predict the longer N-terminal alpha-helical domain of gp63 is in the cytoplasm. The *gp62* gene encodes a 109-amino acid protein with two TMDs and both termini located in the cytoplasmic compartment (shown with an Alphafold3 prediction in Fig. S4) (40, 41). Two alternative start sites with recognizable upstream Shine-Dalgarno sequences for *gp62* are reminiscent of the antiholin dual start motif regulatory module found in the λ S107/S105 gene. Since both N4 *gp62* and *gp63* genes encode alternative start sites, we included their full sequences when cloning the genes to assay their full coding potential.

To investigate the activities of gp62 and gp63, we expressed them individually and together in lambda lysogens encoding spanins with either a soluble endolysin (lacking the lambda S canonical holin, S105^λ^) or a SAR endolysin (lacking the phage 21 pinholin, S^21^) (Fig. 3A-B). Gp63 demonstrated complementation of the S^21^ pinholin defect, but not the S105^λ^ canonical holin defect, suggesting it forms pinholes (Fig. 3A-B). Gp62 did not complement holin activity within the first two hours after induction when other characterized holins usually trigger hole formation (Fig. 3A-B). However, gp62 did exhibit partial complementation after ∼2 hours through gradual lysis of the bulk population in both the S canonical holin and S^21^ pinholin genetic backgrounds, suggesting it forms canonical large holes. Coexpression of gp62 and gp63 in the soluble endolysin background resulted in growth cessation by 30 min as seen for gp63 alone, then asynchronous lysis beginning at 120 min as seen for gp62 alone, confirming gp63 and gp62 both exert independent effects on the cell (Fig. 3A). In contrast, gp62-gp63 coexpression in the SAR endolysin background resulted in an even more rapid decline in OD than either gene expressed alone, suggesting a synergistic effect (Fig. 3B). The complementation assays suggest that gp63 is a pinholin and gp62 is a holin with a late triggering time.

**Figure 3.**
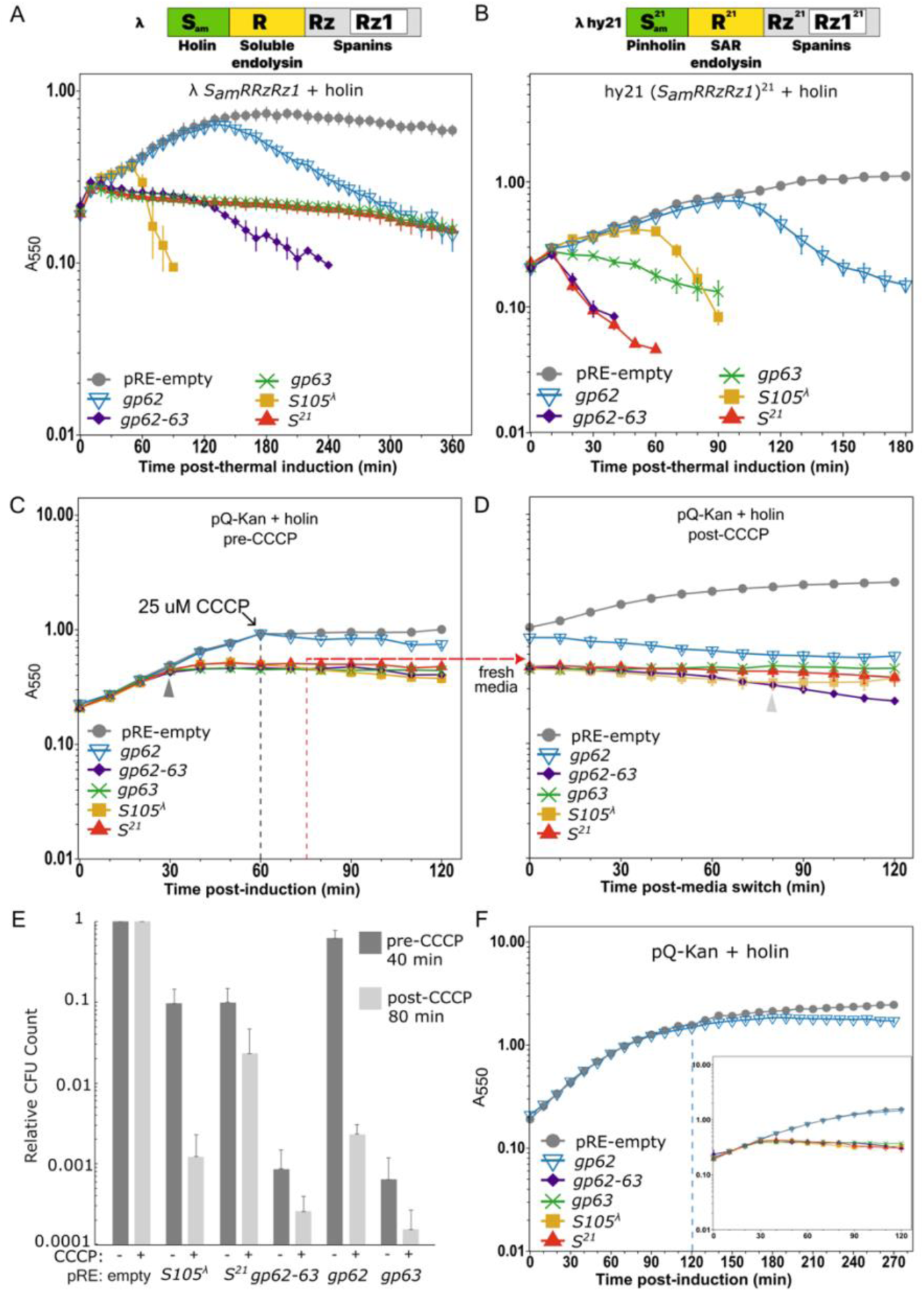
N4 encodes a holin and pinholin. Lysis was monitored in **A)** λ and **B)** hy21 lysogens with amber alleles in the holin as shown in the schematic with simultaneous thermal and IPTG induction. **C)** & **D)** Lysis kinetics of MG1655 cultures induced with IPTG at T=0 minutes with or without 25 µM Carbonyl cyanide m-chlorophenylhydrazone (CCCP) treatment at 60 min (dashed grey line). Culture aliquots were removed, pelleted, and resuspended in fresh media without CCCP at 75 minutes (dashed red line). Strains in A-D carry pRE-empty (solid gray circle), *gp62* (open blue inverted triangle), *gp62*-*63* (solid purple diamond), *gp63* (green x), *S105*^λ^ holin (solid gold square), or *S^21^* pinholin (solid red triangle). **E)** Relative colony-forming units (CFU) for each construct are normalized to the corresponding empty vector control at the respective time point, as marked by arrowheads. **F)** Lysis kinetics of MG1655 cultures with only the plasmids as listed in A-D. Lysis curves and viability counts represent at least three biological replicates and data are presented as the mean ± standard error of the mean.

Due to the late triggering time exhibited by gp62, we used a second assay to demonstrate both proteins exhibit holin activity. A defining characteristic of triggered holins is immediate cell death (15). Treating growing cells with an artificial membrane depolarizer, CCCP, results in temporary growth inhibition until the CCCP is removed (Fig. 3C-D). However, CCCP treatment in cells expressing any type of holin results in irreversible triggering and cell death, even after CCCP removal. When S105^λ^ or S^21^ are expressed, we observe growth cessation and >99% decrease in CFUs (Fig. 3C-E). Gp63 expression alone immediately caused >99% decrease in CFUs and permanent cell death with no recovery after CCCP treatment (Fig. 3C-E). In contrast, gp62 expression alone did not result in significant growth defects (Fig. 3F) until the artificial depolarizer CCCP was added (Fig. 3 C-D), when there was a >98% CFU decrease suggesting irreversible hole formation (Fig. 3E). The cells expressing gp62, gp63, or gp62-63 were unchanged in their rod morphology, similar to vector alone, S105^λ^, and S^21^ cells (data not shown). Together, the CCCP assay and complementations demonstrate that gp63 is a pinholin and gp62 is a canonical large holin with delayed triggering time.

### Mutations in the pinholin and SAR endolysin lead to rapid lysis

Given the presence of a fully functional MGL system, we wondered what regulators prevented rapid lysis during phage N4 infections. Shortly after its initial discovery in the early 1960’s, nitrous acid-induced chemical mutagenesis of N4 led to the identification of the N4 *r* (for rapid lysis) phenotype (7). The original N4 *r* isolate exhibits synchronized culture lysis within an hour, contrasting with the asynchronous WT N4 lysis starting a few hours after infection. We acquired an N4 stock passaged from the early *r* strain. After plaque purification, we sent two lines from that stock for whole genome sequencing (summarized in Table S1). Mutations were observed scattered across the *r* genome, including silent mutations in the vRNAP gene *gp50*, changes in the length of a homopolymeric pyrimidine tract within the direct terminal repeat (DTR), multiple changes in noncoding regions, and missense mutations in middle and late genes. Since the separation between the original WT and *r* lines may have been exaggerated over time due to lab passaging, we could not assign a specific allele to the *r* phenotype with these data.

To build a comprehensive picture of the genetic changes that can lead to rapid lysis, we performed a selection for rapid lysing phage variants. Starting with the parent N4 WT, we collected infection supernatant samples between 20-60 minutes post-infection. Those supernatants yielded mixed plaque phenotypes matching both the smaller WT character and the larger *r* phenotype with fuzzy halos (Fig. 4), akin to the *r* plaques observed in T4 lysis inhibition mutants (19). Over 60 N4 *r* phages were plaque purified. Isolates that lysed within an hour were subjected to targeted sequencing in the lysis cassette. Isolates with no changes in the lysis cassette were whole genome sequenced (summarized in Table 1). We grouped the mutations into four classes: lysis cassette mutants, mostly silent mutants in the vRNAP *gp50* gene, DTR mutants, and miscellaneous changes. All six *r* genotypes measured formed significantly larger plaques than WT (Tukey P ≤ 0.006), but the non-WT genotypes did not significantly differ from one another (all P > 0.75) (Fig. 4A-C). Representatives shown in Fig. 4D display rapid lysis like the original *r* line. Further characterization of the vRNAP and DTR classes will be discussed in a separate report. The miscellaneous class included changes found throughout the genome in both coding and noncoding regions, some co-occurring with the other classes, that were not isolated more than once (Table 1). One of the miscellaneous class missense lines was Q98R in gp53, a late gene whose protein product is predicted to contain TMDs. We built the gp53 Q98R change into a clean N4 WT background, but it did not replicate the rapid lysis phenotype (data not shown). Since these miscellaneous mutations did not predominate or replicate rapid lysis on their own, we analyzed the other classes.

**Figure 4.**
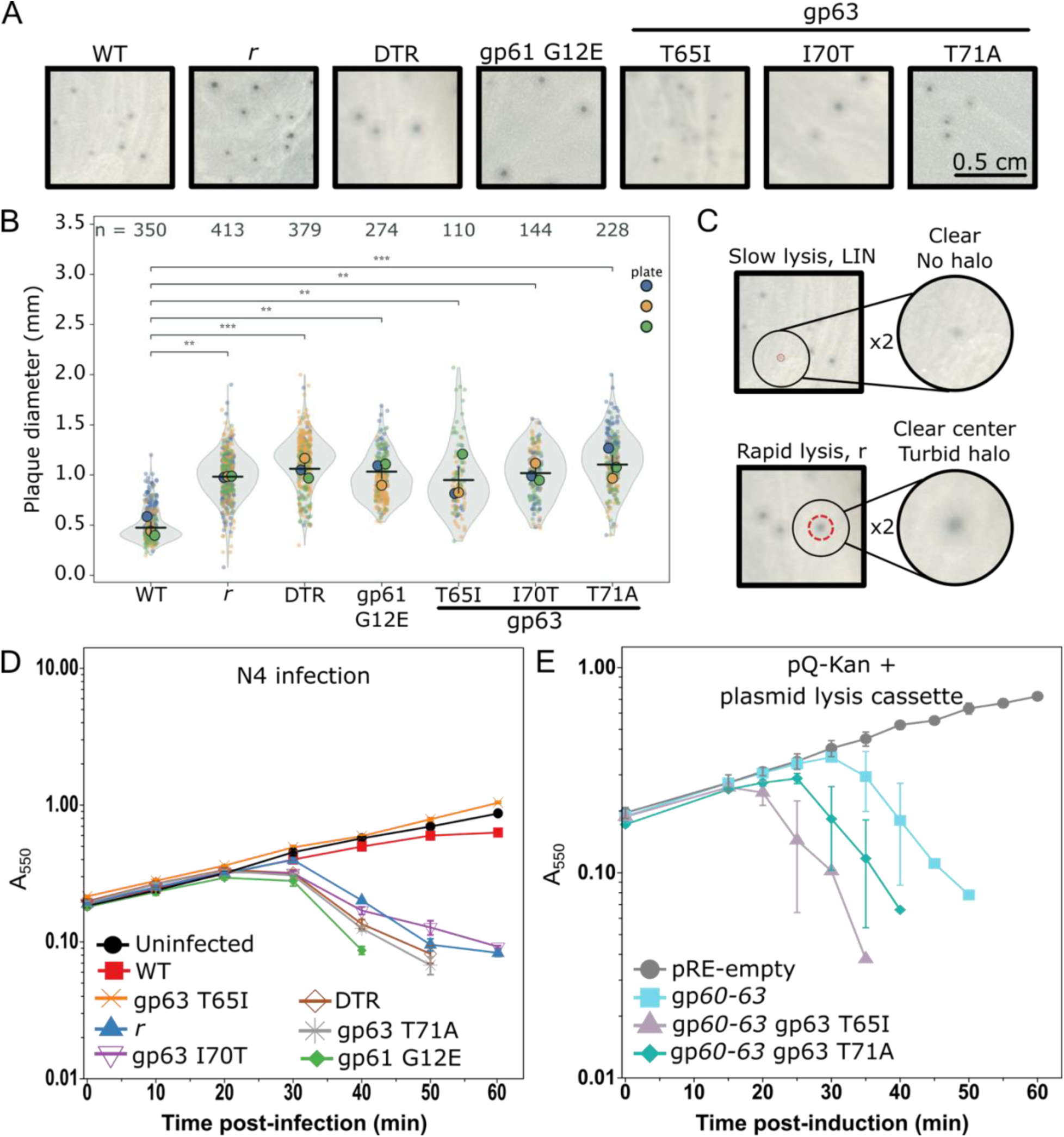
Lysis kinetics of N4 rapid lysers. **A)** N4 plaques for each rapid lyser class compared to the WT parent on host lawns (0.5 cm scale bar). **B)** Full violin distributions for 1,898 total plaque diameters are plotted. Genotypes differed overall by one way ANOVA on plate means; brackets show Tukey post hoc comparisons against WT (* P < 0.05, ** P < 0.01, *** P < 0.001). **C)** Enlarged example plaques for WT and a rapid lyser. **D)** Lysis curve monitoring N4 MOI=5 infections for WT parent (solid red square) and representatives of N4 rapid lyser classes: an original rapid lysis *r* derivative (solid blue triangle), DTR (open brown diamond), gp63 pinholin mutants (orange x, gray asterisk, and open purple inverted triangle), and gp61 SAR endolysin G12E (solid green diamond). **E)** Lysis curve after 1 mM IPTG induction monitoring culture density for pRE-empty vector (solid gray circle), gp*60-63* (solid cyan square), gp*60-63* gp63 T65I (solid pink triangle), or gp*60-63* gp63 T71A (solid green diamond). All lysis curves represent at least three biological replicates and data are presented as the mean ± standard error of the mean.

**Table 1.**
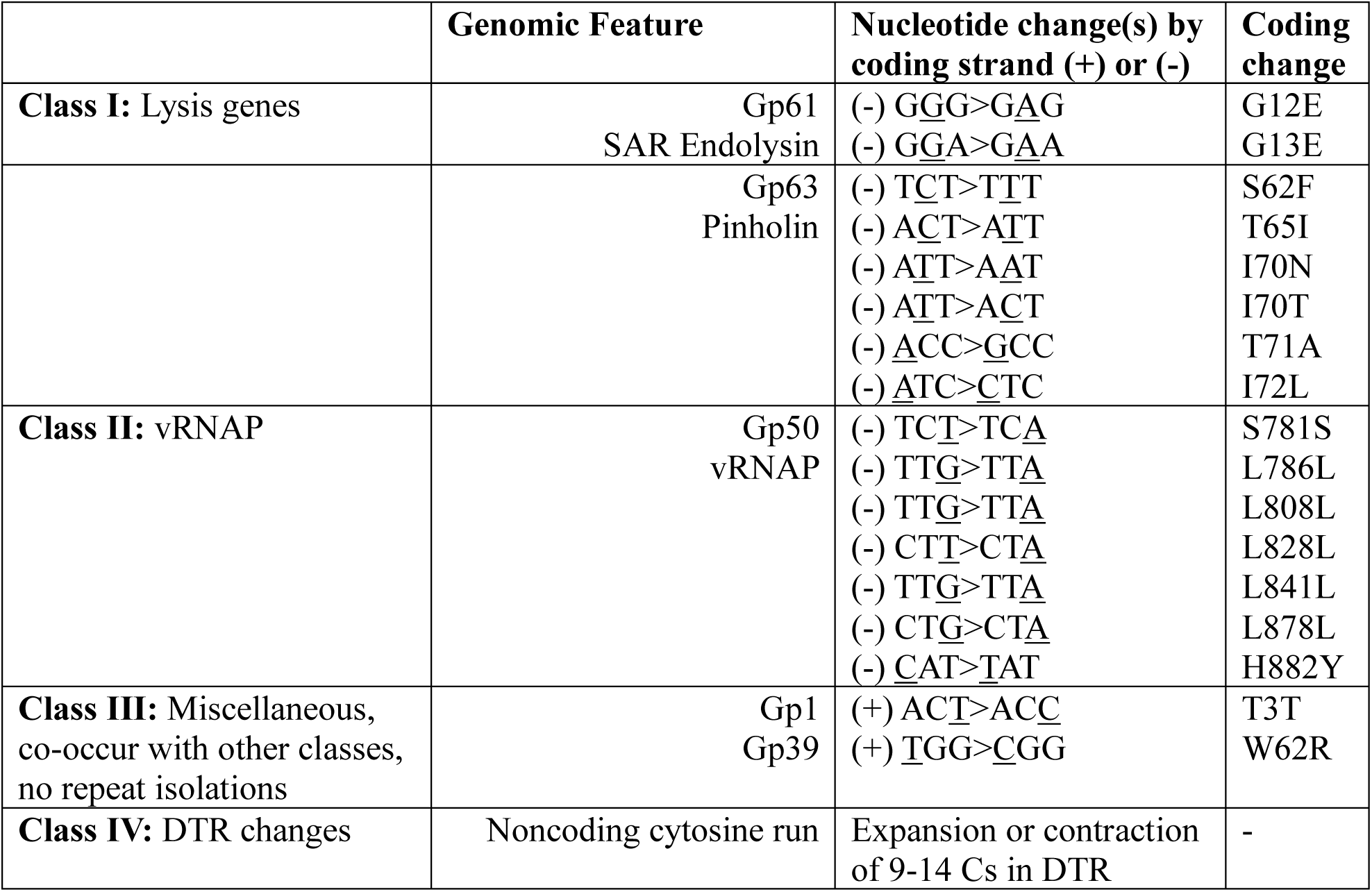
Rapid lysis mutant classes. Summary of spontaneous genetic changes associated with rapid lysis in N4 after selection for early lysis in liquid and/or large *r* plaque phenotype. All nucleotide changes are given relative to the coding strand, which refers to the nontemplate strand of the open-reading frame for a gene, or the plus strand if not a coding region.

|  | Genomic Feature | Nucleotide change(s) by coding strand (+) or (-) | Coding change |
| --- | --- | --- | --- |
| **Class I:** Lysis genes | Gp61<br>SAR Endolysin | (-) <u>G</u>G<u>G</u>><u>G</u><u>A</u><u>G</u><br>(-) <u>G</u><u>G</u><u>A</u>><u>G</u><u>A</u><u>A</u> | G12E<br>G13E |
|  | Gp63<br>Pinholin | (-) <u>T</u><u>C</u><u>T</u>><u>T</u><u>T</u><u>T</u><br>(-) <u>A</u><u>C</u><u>T</u>><u>A</u><u>T</u><u>T</u><br>(-) <u>A</u><u>T</u><u>T</u>><u>A</u><u>A</u><u>T</u><br>(-) <u>A</u><u>T</u><u>T</u>><u>A</u><u>C</u><u>T</u><br>(-) <u>A</u><u>C</u><u>C</u>><u>G</u><u>C</u><u>C</u><br>(-) <u>A</u><u>T</u><u>C</u>><u>C</u><u>T</u><u>C</u> | S62F<br>T65I<br>I70N<br>I70T<br>T71A<br>I72L |
| **Class II:** vRNAP | Gp50<br>vRNAP | (-) <u>T</u><u>C</u><u>T</u>><u>T</u><u>C</u><u>A</u><br>(-) <u>T</u><u>T</u><u>G</u>><u>T</u><u>T</u><u>A</u><br>(-) <u>T</u><u>T</u><u>G</u>><u>T</u><u>T</u><u>A</u><br>(-) <u>C</u><u>T</u><u>T</u>><u>C</u><u>T</u><u>A</u><br>(-) <u>T</u><u>T</u><u>G</u>><u>T</u><u>T</u><u>A</u><br>(-) <u>C</u><u>T</u><u>G</u>><u>C</u><u>T</u><u>A</u><br>(-) <u>C</u><u>A</u><u>T</u>><u>T</u><u>A</u><u>T</u> | S781S<br>L786L<br>L808L<br>L828L<br>L841L<br>L878L<br>H882Y |
| **Class III:** Miscellaneous, co-occur with other classes, no repeat isolations | Gp1<br>Gp39 | (+) <u>A</u><u>C</u><u>T</u>><u>A</u><u>C</u><u>C</u><br>(+) <u>T</u><u>G</u><u>G</u>><u>C</u><u>G</u><u>G</u> | T3T<br>W62R |
| **Class IV:** DTR changes | Noncoding cytosine run | Expansion or contraction of 9-14 Cs in DTR | - |

Here, we focus on the mutations in the lysis proteins gp63 and gp61. The two changes in gp61 introduced a single negatively charged residue into the SAR domain, likely preventing its anchoring in the inner membrane, which would result in rapid peptidoglycan degradation and spanin action leading to rapid lysis (Table 1 and Fig. S2). The six changes identified in gp63 are clustered in a highly conserved region near the TMD (Table 1 and Fig. 5). Five of the six mutations resulted in changes to or from serines and threonines (Fig. 5). The gp63 T65I allele demonstrated variable lysis phenotypes across N4 *r* isolates suggesting reversion or the presence of secondary mutations (Fig. 4D). To verify that these changes in gp63 are involved in rapid lysis, we reconstructed two gp63 alleles into the plasmid lysis system. Both T65I and T71A gp63 alleles demonstrated rapid lysis faster than the parent lysis cassette (Fig. 4E). Due to the clustering of the gp63 changes (Fig. 5), we designated this region the LIN domain and postulate it serves a regulatory function. Our screening of over 60 individual plaques did not isolate a single gp62 mutant, suggesting gp62 is not responsible for establishing LIN. The multiple SAR endolysin and pinholin alleles that result in rapid lysis suggest they are dominant factors controlling lysis timing in phage N4.

**Figure 5.**
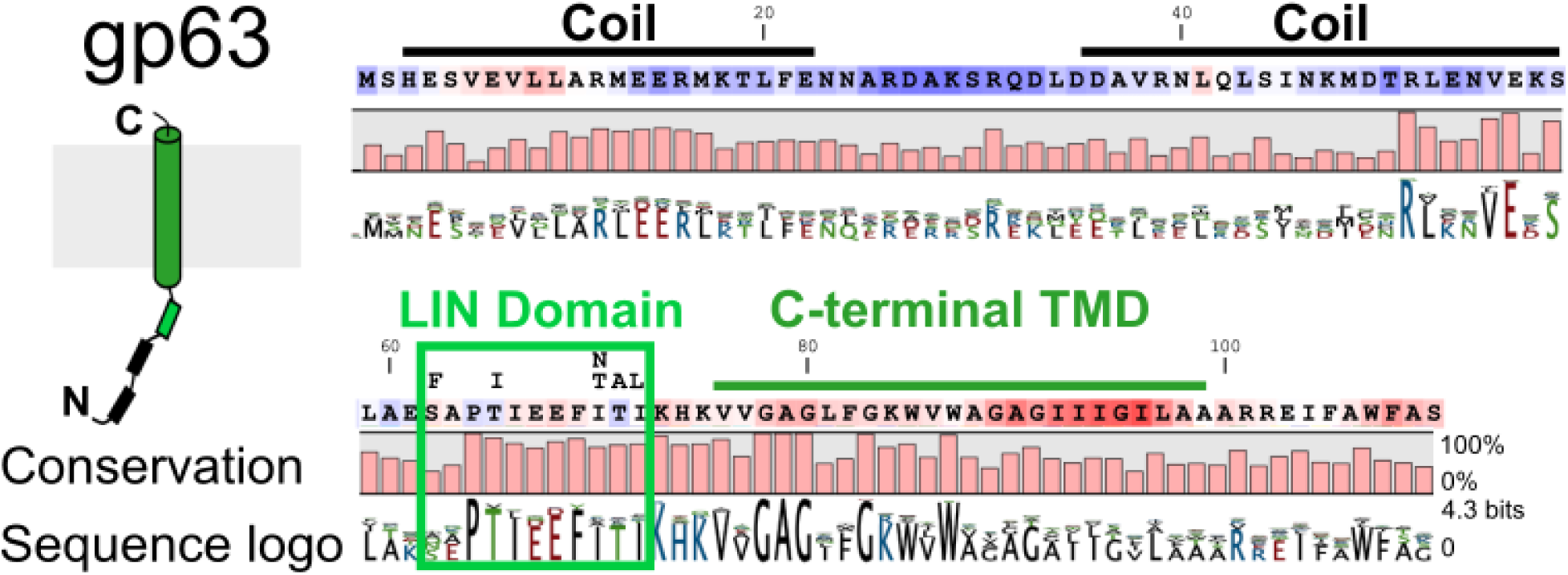
Gp63 pinholin conservation. N4 gp63 protein sequence shown in Kyte-Doolittle hydropathy color scale where blue is hydrophilic and red is hydrophobic. Predicted coiled-coil regions and the C-terminal anchor transmembrane domain (TMD) are delineated. The LIN domain outlines the cluster of mutations found associated with rapid lysis: S62F, T65I, I70T, I70N, T71A, and I72L. The conservation plot and sequence logo summarize a multiple sequence alignment of 80 proteins from phages and bacterial hits with significant sequence similarity to gp63 by BLASTp and BLASTx (full alignment in Fig. S6).

### The pinholin, but not the holin, are essential for lysis

We further explored a role for gp62 in lysis during phage infection. When we expressed excess gp62 from a plasmid in both WT and *r* infections, we observed no change in lysis timing or plaque morphology (data not shown). Next, we deleted *gp62* in phage N4 in both WT and rapid lysis isogenic backgrounds. The Δ62 phage plaques were identical to the parent phage (Fig. S5A). The lysis profile for infections with N4 *r* parent and its isogenic Δ62 phage were identical (Fig. S5B). In contrast, the long lysis delay in N4 WT was significantly shortened in its isogenic Δ62 phage, with lysis onset an hour earlier and the rate OD decrease enhanced. However, gp62 expression from a plasmid did not complement the lysis timing (Fig. S5C), preventing us from concluding more than that gp62 is nonessential from the WT Δ62 phage.

Since both gp62 and gp63 exhibited holin activity, we hypothesized that gp62 could substitute for gp63 during phage lysis. To test this, we deleted *gp63* from the N4 genome and recovered viable phage in both the WT and *r* backgrounds (Fig. S5A). Rapid lysis was observed in the WT Δ63 background (Fig. S5D), however, sequencing revealed that the recovered recombinant phages encoded secondary mutations in gp61 that we demonstrated above lead to rapid lysis (Table S2). Additionally, expression of gp63 from a plasmid did not recover the slow lysis phenotype in the WT Δ63 lines with different secondary mutations (Fig. S5E). Deletion of gp63 from the N4 *r* phage genetic background had no impact on lysis timing (Fig. S5F-G). Therefore, we conclude that while gp62 is not essential for lysis or LIN, gp63 is conditionally essential for lysis.

### Conservation of the N4 lysis proteins

Among all the 144 high-quality cultured N4-like virus contigs identified with at least seven core genes (33), the lysis proteins were not well annotated, making their conservation unclear. Using BLASTp and BLASTx we searched for bacterial or phage proteins with sequence similarity to gp63 or gp62. The hits were distributed across bacterial assemblies and N4-like phages, plus one plasmid contig. Many of the bacterial contigs, like the plasmid, were approximately the size of a complete N4 genome suggesting they were complete phage sequences where a functional lysis cassette would be present. For phages with BLAST hits to only gp62 or gp63, manual genome inspection usually revealed a misannotated or unannotated pair protein that we added to our list. Gp62 and gp63 co-occurred in 60/65 N4-like genomes with hits (Table S3). After deduplication to remove identical sequences, 80 gp63-like proteins showed a variable N-terminal cytoplasmic tail followed by highly conserved transmembrane and LIN domains (conservation plot in Fig. 5, complete alignment in Fig. S6). The deduplicated protein alignments for gp62-like proteins showed conservation of the two TMDs with a few residues conserved in nearly 100% of the 86 aligned proteins (conservation plot in Fig. 6, complete alignment in Fig. S7). Their co-occurrence and conserved transmembrane and LIN regions are consistent with a regulatory role for gp62 in gp63-mediated lysis events.

**Figure 6.**
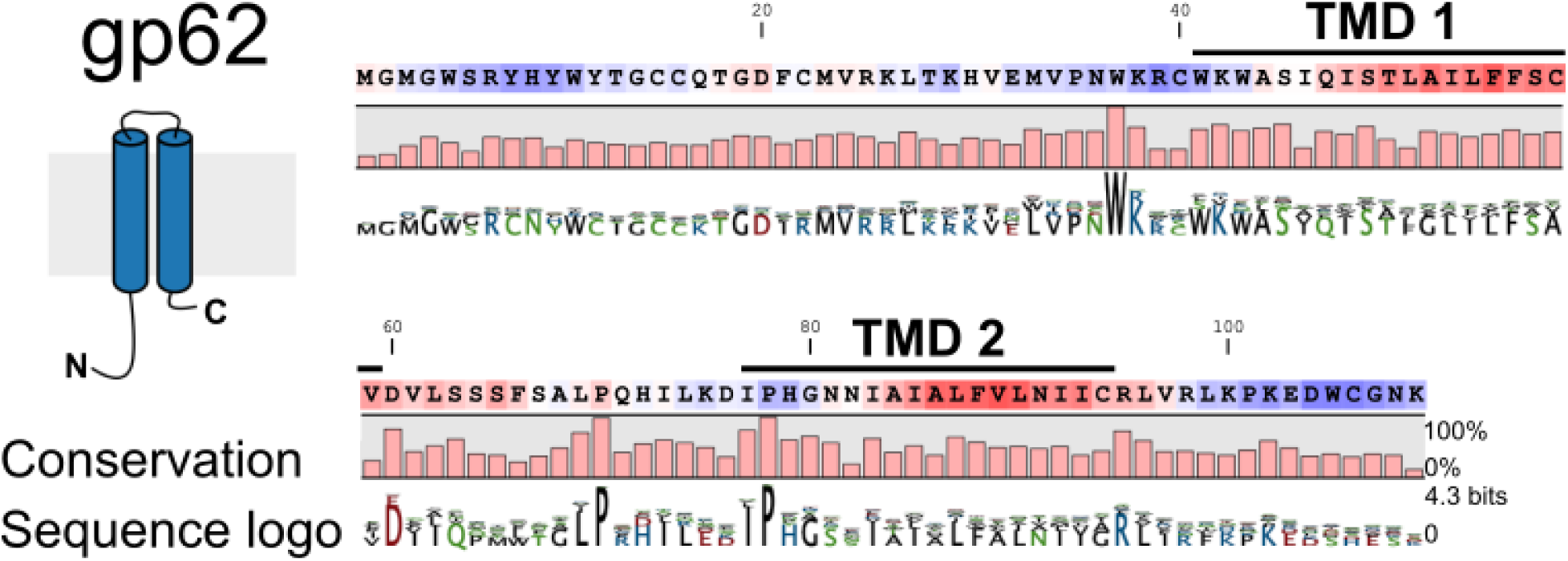
Gp62 holin protein conservation. N4 gp62 protein sequence shown in Kyte-Doolittle hydropathy color scale where blue is hydrophilic and red is hydrophobic. Predicted transmembrane domains (TMD) are delineated. The conservation plot and sequence logo summarize a multiple sequence alignment of 86 proteins from phages and bacterial hits with significant sequence similarity to gp62 by BLASTp and BLASTx (full alignment in Fig. S7).

## Discussion

Here, we demonstrate that phage N4 has a fully functional MGL lysis system composed of a pinholin (gp63), holin (gp62), SAR endolysin (gp61), and bimolecular spanins (gp60 and gp60a). Though WT phage N4 exhibits LIN with a long asynchronous lysis, expression of the N4 MGL system from a plasmid effects rapid, synchronous culture lysis. By selecting for phage exhibiting rapid lysis, we isolated spontaneous rapid variant alleles in the gp63 pinholin and gp61 endolysin genes, suggesting that N4 gp63 pinholin inhibition produces the LIN state. We propose that phage N4 exits the LIN state by gp62 holin activity promoting release of the SAR endolysin (Fig. 7).

**Figure 7.**
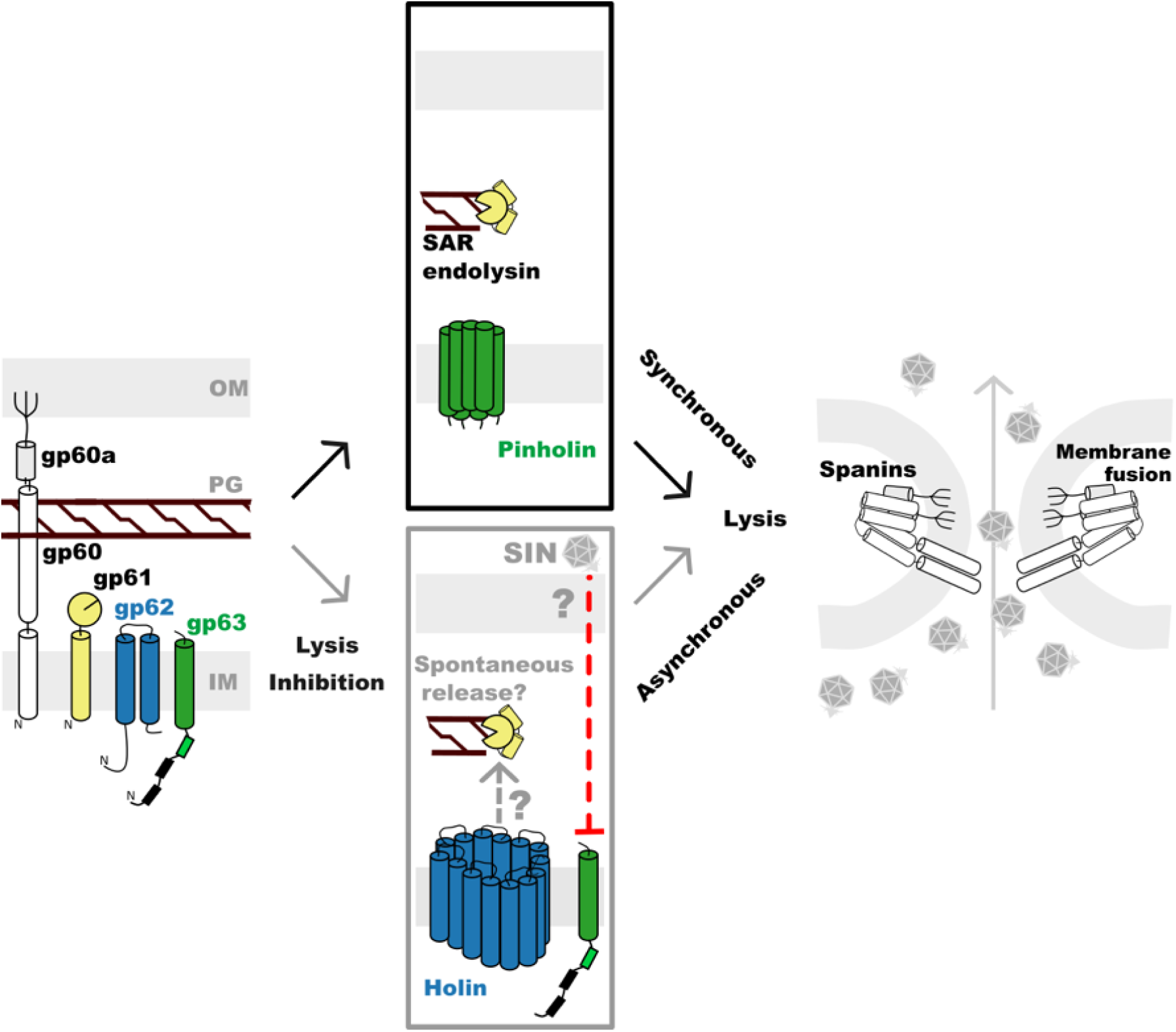
N4 lysis model. Phage N4 lysis proteins are the spanins (gp60 and gp60a), SAR endolysin (gp61), holin (gp62), and pinholin (gp63). The gp63 pinholin can effectuate release of the gp61 SAR endolysin to cause the rapid and synchronous lysis phenotype, as seen in *r* phage. In the presence of superinfection (SIN), lysis inhibition (LIN) results. LIN seems to be due to inhibition of the gp63 pinholin. Gp61 SAR endolysin activity, possibly released spontaneously or by the late gp62 holin, leads to asynchronous lysis and phage release from cells in the LIN state. Black and colored text are demonstrated aspects of lysis and LIN. Gray text in the LIN pathway indicates a mechanistic model proposed here.

Under the canonical pinholin-SAR endolysin lysis model, rapid lysis results from holin or pinholin holes synchronously releasing the SAR endolysin. We show phage N4 uses a gp63 pinholin-gp61 SAR endolysin system. Consistent with the stochastic leakiness of SAR endolysins, we show that low levels of gp61 induction alone cause loss of cell wall integrity (Fig. 2C). Co-expression of the spanins with low levels of gp61 leads to rapid cell lysis. Stojković and Rothman-Denes previously demonstrated that overexpression of gp61 alone, which would have a higher level of leaky SAR endolysin, leads to rapid and complete cell lysis (27, 42). During the N4 LIN state, when gp63 pinholin activity is inhibited, stochastic SAR endolysin release or gp62 holin-mediated synchronous release could lead to rapid individual cell lysis that manifests as a gradual lysis in the bulk OD population measurement (18, 43). Moreover, gp63 deletion required secondary mutations in the endolysin SAR domain such that early release would be inevitable, cementing the link between SAR endolysin release and the rapid lysis phenotype (Fig. S5). Prior studies also suggested that gp63 is an activator of gp61 (Stojković, 2006 PhD thesis) (27, 42). Overall, these data support a major role for the SAR endolysin in both typical lysis and exit from LIN.

Unlike the typical sharp drop in OD observed during rapid lysis, the exit from LIN manifests as asynchronous lysis. Our data support the conclusion that gp63 is the holin functioning in rapid lysis and gp62 is the holin that supports exit from LIN. Since the asynchronous lysis pattern is mimicked by the gp62 holin when it is the only holin in a cell (Fig. 3), we propose that lethal activity from accumulated gp62 promotes gp61 release. Indeed, the phenotypes displayed by gp62 are consistent with multiple lysis regulation activities. First, gp62 expressed from a plasmid is lethal to the cell (Fig. 3), suggesting gp62 forms irreversible holes. Although gp62 lethality manifests more than an hour after an uninhibited gp63 pinholin triggers, its activity coincides with the time when N4 WT exits from LIN via asynchronous lysis (Fig. S5). However, expression differences between our constructs may not reflect the precise ratios found during phage infection, therefore this should be accounted for in future studies. Second, gp62 expression promotes the release of both the phage 21 SAR endolysin and the soluble λ endolysin, a globular enzyme of 17.8 kDa, across the inner membrane (Fig. 3). Third, gp62 is nonessential to the phage as it can be deleted without impairing lysis (Fig. S5). Finally, although our bioinformatics showed that most phages encode gp62 and gp63 as a pair, this does not preclude an independent gp62 activity in LIN exit. Structural modeling of gp62-gp63 multimers using Alphafold2 or 3 does not reveal significant interactions (data not shown), likely a limitation of the prediction algorithms that do not restrict orientation for proteins normally bound in a membrane environment. Altogether, these data suggest gp62 is a nonessential holin that promotes lysis during exit from LIN, whereas gp63 promotes rapid, synchronous lysis.

Despite their genotypic differences, the LIN phenotypes displayed by phages N4 and T4 operate on common principles. Broadly, both phages exhibit delayed lysis timing in response to high phage density (7, 19). The LIN state in both N4 and T4 lasts for hours, leading to significantly increased burst size. Both N4 and T4 rapid lysis variants manifest with larger plaques that have mutations in regulatory proteins or the holin gene itself. A key lynchpin we report here is that holin regulation is essential for LIN in N4 as well as T4. However, the molecular pathways that regulate the holin in T4 and N4 are likely completely distinct. Beyond the lack of amino acid identity between their lysis and known regulatory proteins, their subcellular localizations are totally different. T4 has soluble antiholins rIII in the cytoplasm and rI in the periplasm, as well as the periplasmic spackle that functions in superinfection exclusion (44, 45). The periplasmic T4 proteins are essential for the inhibitory interaction with the T holin (22, 46). In contrast, all the N4mutations associated with rapid lysis thus far are in membrane proteins, including the membrane-anchored SAR endolysin and the cytoplasmic region of the gp63 pinholin. The simplest explanation for these differences in protein localization is that N4 LIN is not triggered by regulatory periplasmic interactions like T4 (47), but this remains to be rigorously tested. Interestingly, LIN is documented in the distant T4-like *Vibrio cholerae* phage ICP1 and *Salmonella* phages Manna and Mansal (26, 48). In N4-like phages, the *E. coli* phage G7C does not exhibit LIN, while *Shigella* N4-like phages may (34, 49). Therefore, the extent of mechanistic conservation and distinction among lysis and LIN pathways within these two major phage types remains to be explored.

With the primary N4 lysis proteins identified, the other rapid lysis mutant classes reported here will be mined for inhibitory factors that trigger LIN. Studying the relationship between N4 lysis and LIN will reveal the mechanistic underpinnings connecting lysis timing and yield in a phage with an average burst size of ∼3000 virions/cell after a few hours compared to the typical ∼100-300 virions/cell in an hour (50, 51). Future comparisons among diverse N4-like phages could reveal common and unique lysis regulation principles that govern phage yield.

## Materials and methods

### Bacterial strains and growth conditions

All *E. coli* MG1655 derivatives (Table 2) and phages (Table 3) used in this study are listed in the tables. Cultures were grown in lysogeny broth (LB) medium, composed of 10 g/L tryptone, 5 g/L yeast extract, and 10 g/L NaCl (no magnesium supplementation) with aeration at 37°C unless otherwise stated, except lysogens which were maintained at 30°C until thermal induction. Solid medium for plates was prepared by adding 1.5% (w/v) agar. For phage plaque assays, a soft agar overlay consisting of LB with 0.75% (w/v) agar was used. Strains harboring plasmids were maintained under appropriate antibiotic selection: ampicillin at 100 µg/mL, kanamycin at 40 µg/mL, or chloramphenicol at 10 µg /mL. Glucose at 0.4% (w/v) was added to repress leaky lysis protein expression in their respective maintenance cultures.

**Table 2.**
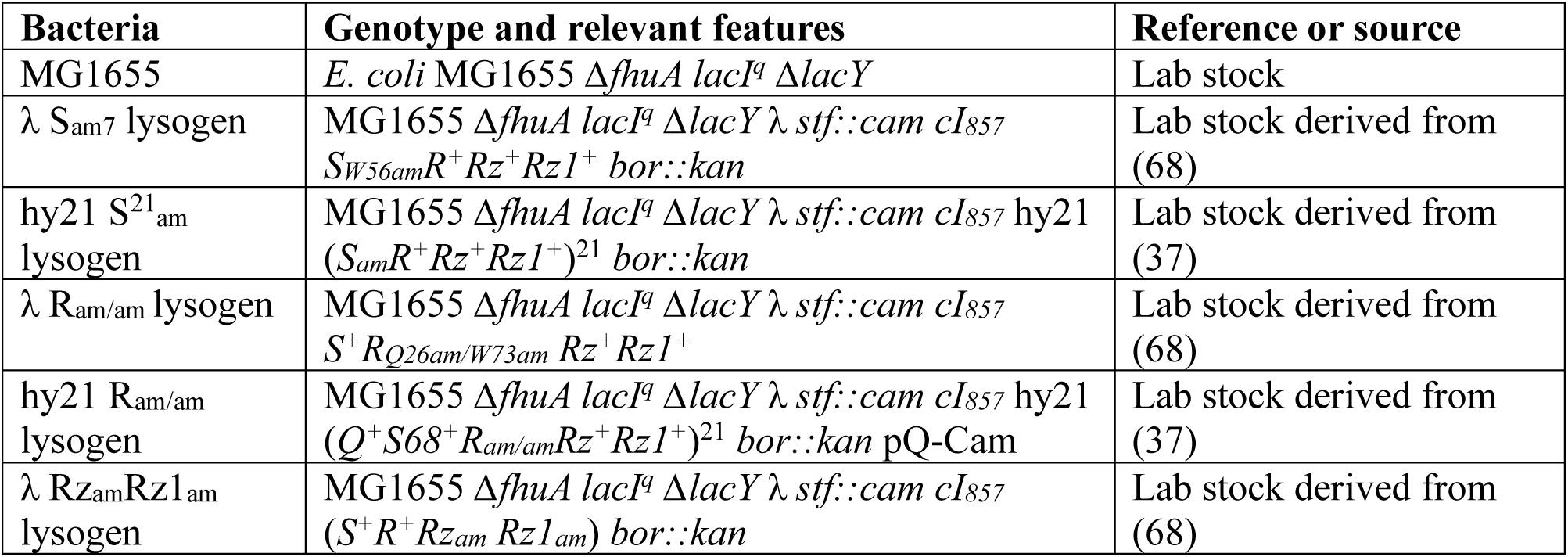
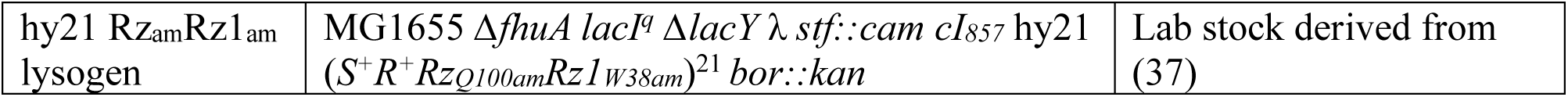
Bacteria used in this study.

**Table 3.** Phages used in this study.

| Phage | Genotype and relevant features | Reference or source |
| --- | --- | --- |
| N4 WT | Virulent *E. coli* K-12 podophage, slow lyser parent | Gift from Ian Molineux (69) |
| N4 *r* | N4 *r* isolated based on plaque size after nitroguanosine mutagenesis, does rapid lysis | Gift from Ian Molineux (7) |
| N4 WT $\Delta 62$ | WT derivative, 215 bp deletion of gp62 residues T28-L99, derived from N4 WT | This study |
| N4 *r* $\Delta 62$ | N4 *r* derivative, 215 bp deletion of gp62 residues T28-L99, derived from N4 *r* Class II/III | This study |
| N4 WT $\Delta 63$ | WT derivative, 247 bp deletion from 14 bp 3' of *gp64* terminator through *gp62*, derived from N4 WT | This study |
| N4 *r* $\Delta 63$ | N4 *r* derivative, 247 bp deletion from 14 bp 3' of *gp64* terminator through *gp62*, derived from N4 *r* Class II/III | This study |

For infections and inductions, a saturated host culture was subcultured at a 1:200 dilution in 25 mL of LB and grown to a mid-log absorbance at 550 nm (A_550_ or OD) of approximately 0.2, then induced or infected. For phage infections, mid-log cultures were incubated on ice during a 10-minute adsorption. Unless otherwise stated, infections were performed at a multiplicity of infection (MOI) of 5. For thermal induction, lysogens were grown with aeration at 30°C, shifted to 42°C for 15 minutes to induce the prophage, then transferred to 37°C. For chemical induction of plasmid-borne phage genes, 1 mM isopropyl-β-D-thiogalactopyranoside (IPTG) was added. λ lysogens were only thermally induced as the Q antiterminator produced from the phage genome will antiterminate the pR’ transcripts from the pRE vector. Hy21 lysogens encode the phage 21 Q, therefore they were paired with the pQ plasmid carrying λ Q and induced both thermally and with IPTG to antiterminate transcripts of the pRE vector.

### Construction of plasmids and phages

Individual N4 genes and combinations of genes were cloned to generate plasmids listed in Table 4 using oligos listed in Table 5. Clones were verified by Sanger sequencing with Eton Bioscience Inc (San Diego, CA) or Oxford Nanopore long-read sequencing with Plasmidsaurus Inc. (Louisville, KY). The predicted lysis cassette of phage N4 was PCR amplified and cloned into restriction sites BamHI and KpnI in the pRE vector by ligation. To generate the gp63 alleles associated with rapid lysis in the pRE plasmid lysis cassette, the full lysis cassette plasmids were amplified with primers containing the base changes to generate T65I and T71A by overlap PCR. The λ R clone in pRE had an mCherry fusion removed via omission PCR followed by ligation.

**Table 4.** Plasmids used in this study.

| Plasmids | Genotype and relevant features | Reference or source |
| --- | --- | --- |
| pQ | $\lambda$ Q cloned under P<sub>lac/ara-1</sub> promoter in a low copy number plasmid pZS-24 | (70) |
| pQ-Cam | pQ plasmid encoding chloramphenicol acetyltransferase | (70) |
| pQ-Kan | pQ plasmid encoding aminoglycoside phosphotransferase from Tn5 | (70) |
| pRE-empty | Medium copy plasmid containing Q-dependent $\lambda$ late promoter pR' | (39) |
| pRE N4 gp60 | N4 genes *gp60* and embedded *gp60a* | This study |
| pRE N4 gp61 | N4 gene *gp61* | This study |
| pRE N4 gp62 | N4 gene *gp62* | This study |
| pRE N4 gp63 | N4 gene *gp63* | This study |
| pRE N4 gp62-63 | N4 genes *gp62* & *gp63* | This study |
| pRE N4 gp60-61 | N4 genes *gp60*, *gp60a*, & *gp61* | This study |
| pRE N4 gp60-63 | N4 genes *gp60*, *gp60a*, *gp61*, *gp62*, & *gp63* (lysis cassette 1678 bp) | This study |
| pRE N4 gp60-62Δ63 | N4 genes *gp60*, *gp60a*, *gp61* & *gp62* (lysis cassette 1427 bp) | This study |
| pRE N4 gp60-63Δ62 | N4 genes *gp60*, *gp60a*, *gp61*, & *gp63*, 215 bp deletion of gp62 residues T28-L99 | This study |
| pRE N4 gp60-63 gp63 T65I | Derivative of pRE N4 gp60-63 | This study |
| pRE N4 gp60-63 gp63 T71A | Derivative of pRE N4 gp60-63 | This study |
| pRE R | R<sup>λ</sup> | This study |
| pRE R<sup>21</sup> | phage 21 R<sup>21</sup> | This study |
| pRE RzRz1 | λ RzRz1 (96% & 98% identity to phage 21 RzRz1) | (71) |
| pRE S105<sup>λ</sup> | λ S105-sfGFP | (72) |
| pRE S<sup>21</sup> | phage 21 S<sup>21</sup>-sfGFP | Lab stock |
| pBA560 | Encodes eLbuCas13a | Gift from Jennifer Doudna, Addgene plasmid # 186236; http://n2t.net/addgene:186236; RRID:Addgene_186236 (52) |
| pBA560-Cas13a-gp62 | Encodes eLbuCas13a and spacer targeting gp62 | This study |
| pBA560-Cas13a-gp63 | Encodes eLbuCas13a and spacer targeting gp63 | This study |
| pTR_sJExD-rfp | Crystal violet-inducible *rfp* under P<sub>JExD</sub> | Gift from Michael Thelen (73) |
| pTR_sJExD | *rfp* deleted from pTR_sJExD-rfp | This study |
| pTR_sJExD - Δ63 | pTR_sJExD with 500 bp of N4 late gene region with the 247 bp of gp63 gene not overlapping with gp62 or its RBS deleted, after gp64 terminator | This study |

**Table 5.**
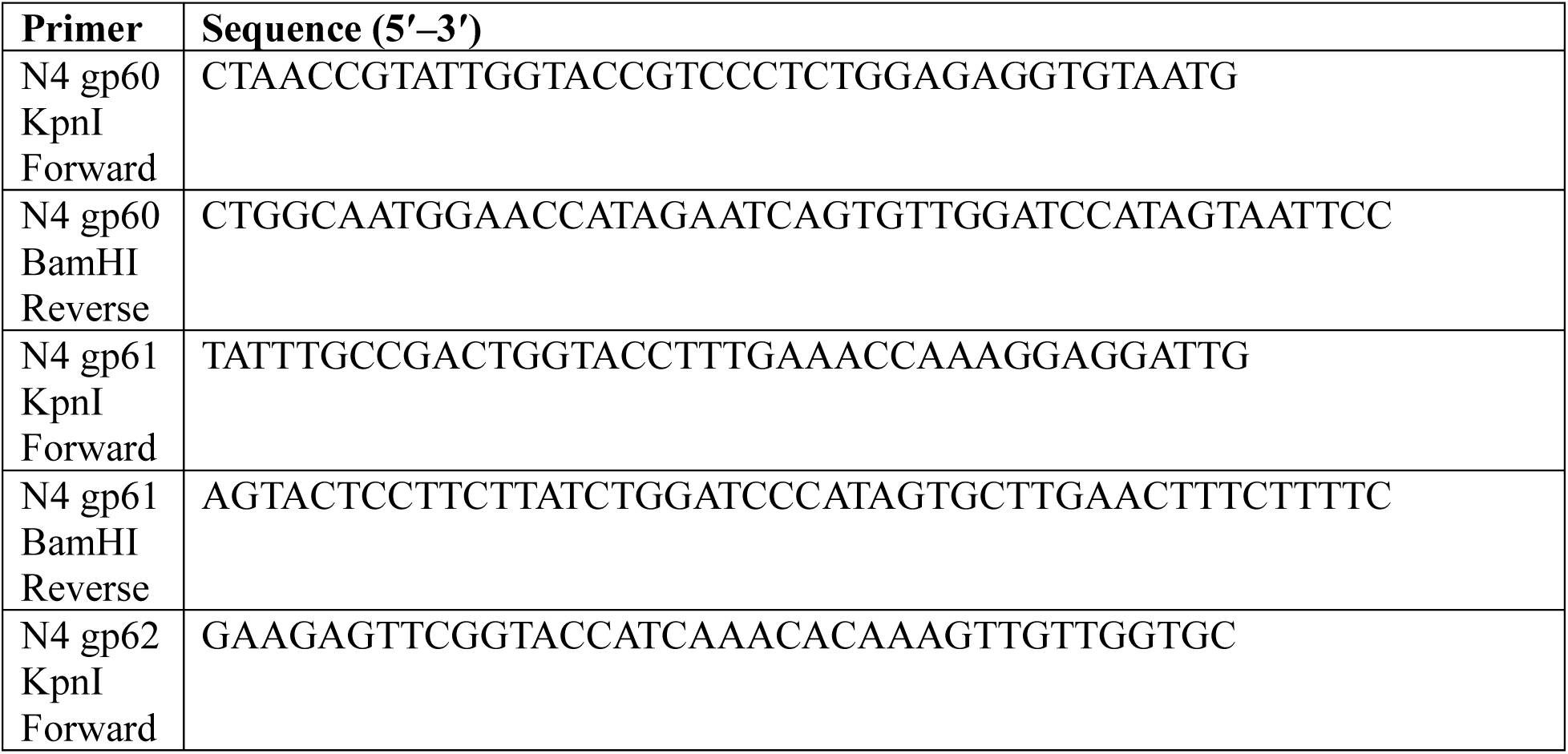
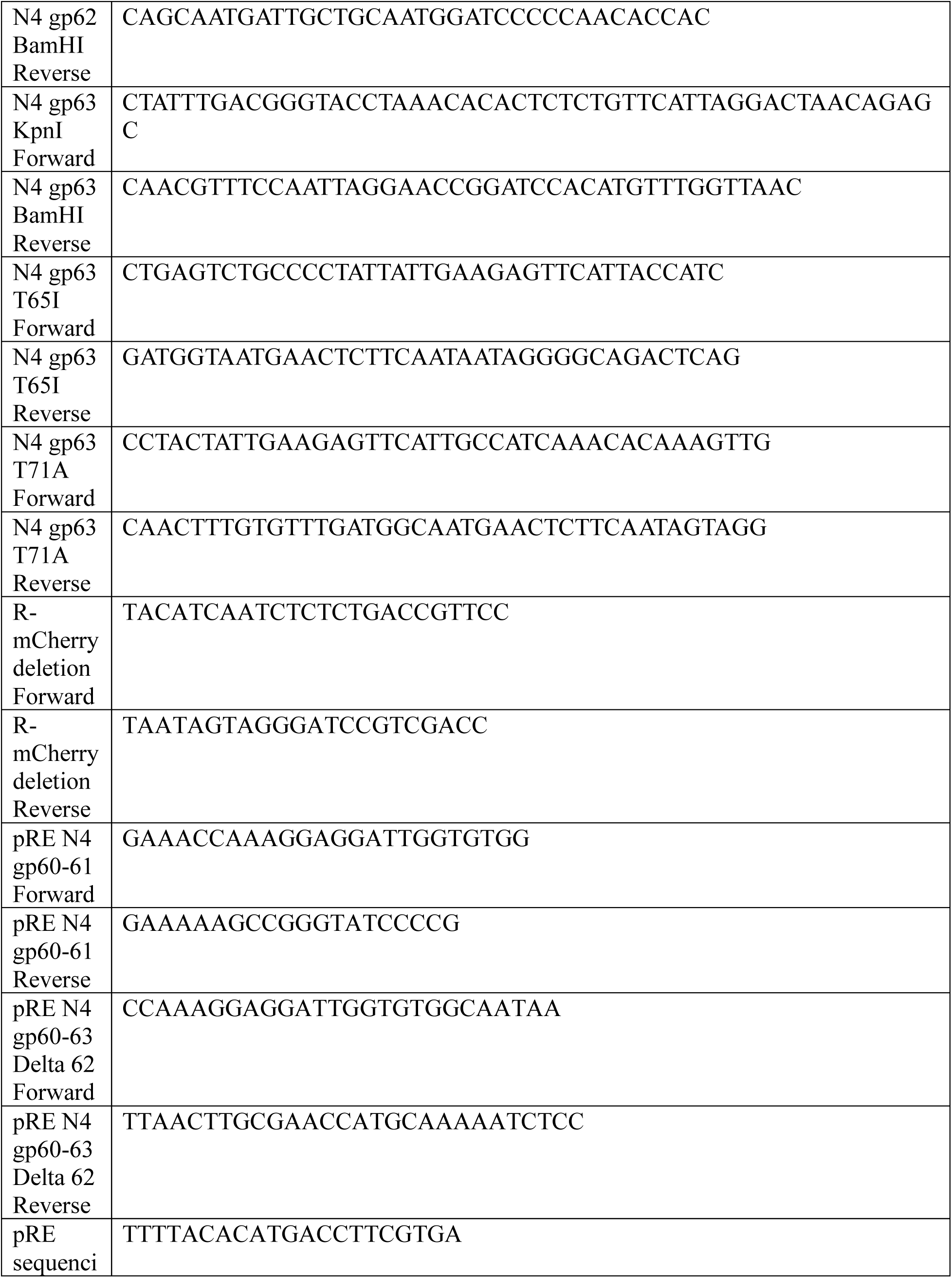
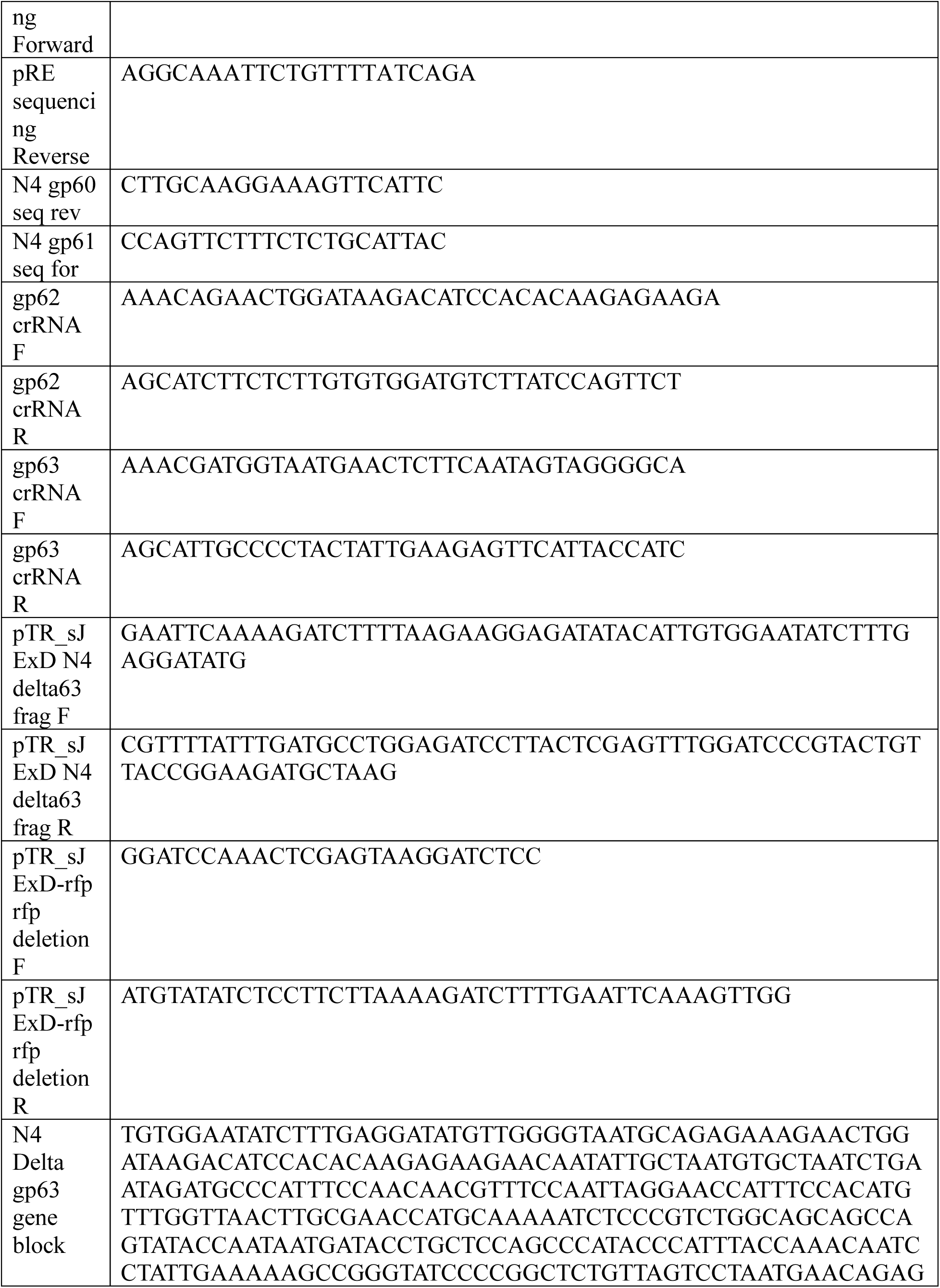
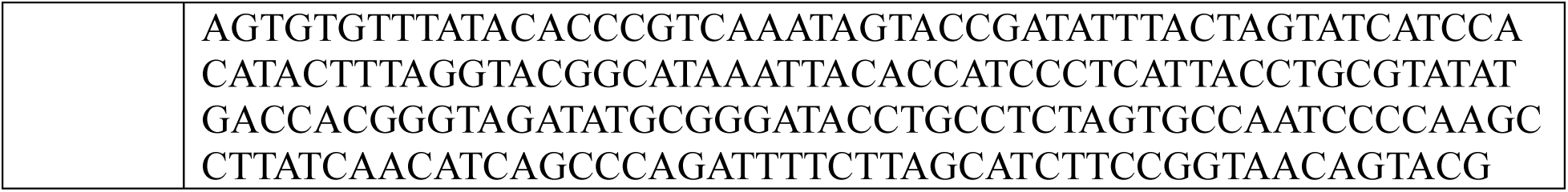
Oligonucleotide sequences and synthesized gene blocks used in this study.

The N4 gp62 and gp63 deletions was constructed by infection of host cells containing a plasmid with the deletion at an MOI=5, 0.1, or 0.01 (all yielded positive recombinants). The Δgp63 knockout plasmid was generated by Gibson assembly of a synthetic gene block from Twist Biosciences into the pTR_sJExD vector. Cross-lysates with recombinant phage were collected and sterilized after an overnight incubation. The parent phages in cross-lysates were counterselected against on lawns with pBA560 encoding eLbuCas13a and a spacer targeting gp62 or gp63 with 0.5-0.6 uM anhydrotetracycline over at least three rounds of plaque purification, according to the method of Adler *et al.*(52). Whole genome sequences were verified using Illumina services at Microbial Genome Sequencing service, now SeqCenter LLC (Pittsburgh, PA), or Plasmidsaurus Inc. (Louisville, KY) Oxford Nanopore Technology long-read sequencing services.

### Plaque morphology and imaging

To visualize plaque morphology, 100 µL of phage dilution was mixed with 100 µL of a saturated overnight host culture. The mixture was incubated on ice for 10 minutes to facilitate phage adsorption. Subsequently, the phage-bacteria mixture was added to 4 mL of molten soft agar and immediately poured onto an LB agar base plate. Plates were incubated overnight at 37°C until plaques developed. Representative plaques were then photographed using an iPhone 14 Pro Max camera set to 2.5X Portrait Mode.

Plaque diameters were measured using Plaque Toolkit v1.0.6 (https://github.com/mbaffour/plaque-toolkit), a desktop application built with the assistance of an AI-coding assistant on the PlaqSeg v0.2.1 segmentation model as the deep-learning detector and the adaptive-local plaque-detection algorithm of Plaque Size Tool (53, 54). Plaque Toolkit measurements were validated against manual measurements made in FIJI/ImageJ (55) with a Pearson correlation coefficient r = 0.979 with R² = 0.958 and p < 0.001. Per plate mean plaque diameters were used for comparisons (n = 3 plates per genotype), following the SuperPlots framework for nested data (56). Homogeneity of variance was confirmed by Levene’s test (P = 0.87). Because replicate numbers were small (n = 3), a parametric test was applied to the plate means rather than an underpowered nonparametric alternative. Genotypes were compared by one way ANOVA (F₆,₁₄ = 8.41, p = 5.4 × 10⁻⁴; effect size η² = 0.78), and pairwise differences were assessed by Tukey’s honestly significant difference test, with adjusted p values reported. Analyses and plots were generated in Python 3.10 using SciPy 1.15.3 (scipy.stats.f_oneway, scipy.stats.tukey_hsd), NumPy 1.26.4, pandas 2.3.3, and Matplotlib 3.10.9.

### Microscopy

Phage-infected cells were placed directly from a shaking culture onto a glass slide under a coverslip. Time series observations were acquired with an Axiocam 702 mono camera mounted on a Zeiss Axio Observer 7 inverted microscope using the alpha plan-apochromat 100×/1.46 oil (UV) Ph3 oil M27 objective. Exported videos were processed using the Carl Zeiss Zen Blue 2.3 imaging software.

Cell morphology after infection or endolysin expression was observed after fixation in 2% glutaraldehyde and 8.6% formaldehyde and Phosphate-Buffered Saline washing. Fixed cells were mounted directly onto glass slides under coverslips and imaged at 40× magnification. Cell detection and measurement extraction used MicrobeJ in Fiji with default cell exclusion presets followed by manual visual inspection to remove artifacts and incorrectly detected objects (57). Morphometric distributions were plotted in R v4.3.2 using ggplot2, with data processing performed using dplyr.

### Rapid lysis phenotype selection and genotyping

Spontaneous mutations leading to rapid lysis were selected in two ways. Initially, spontaneous larger plaques with halos that resemble *r* plaques were plaque-purified directly from plates of WT phage. This method was low frequency and not predictable, but isolates that lysed within an hour were obtained by this method. To select for genotypes associated with rapid lysis, infected culture supernatant was collected at 20-, 30-, or 60-minutes post-infection to enrich for early alleles. This was repeated up to three times followed by plaque purification.

Targeted sequencing was performed by Sanger sequencing of lysis cassette PCR amplicons using the oligos listed in Table 5. For isolates that lysed within an hour but had no mutations in the lysis cassette, phage genomic DNA was extracted from PEG-precipitated capsids and sequenced by Illumina as described previously (58). Assembly and annotation were completed in the Center for Phage Technology Galaxy bioinformatics suite followed by consensus analysis in SnapGene v8.1 (59). Reads were mapped and analyzed for differences using Snippy at usegalaxy.eu (60).

### Holin-CCCP assays

Fresh overnight cultures were diluted, grown, and induced as described above. Sixty minutes post IPTG-induction, holins were triggered by the addition of Carbonyl cyanide m-chlorophenylhydrazone (CCCP) to a final concentration of 25 μM. Fifteen minutes after CCCP addition, 15 mL of the culture was pelleted, the supernatant was removed, and the cell pellet was resuspended in 15 mL of pre-warmed LB containing 1 mM IPTG. This culture was then aerated and monitored simultaneous with the paired treatment flasks.

To assess bacterial viability before and after CCCP treatment, samples were collected at two time points: 40 minutes post-induction (prior to CCCP addition) and 80 minutes after induction in cultures where CCCP had been removed. At each time point, 100 ul of culture was serially diluted in fresh LB and plated in triplicate. Colonies arising after overnight incubation at 37°C were counted and expressed as relative CFU counts compared to the matched pRE-empty control collected at the same time point.

### Bioinformatic analyses of protein sequences

Similar protein sequences were identified through BLASTx and BLASTp searches with the gp62 109 amino acid sequence and gp63 110 amino acid sequence against the non-redundant protein sequences database (retrieved May 4, 2025) and clustered using CD-HIT for 100% identity clusters (61–63). Phages without both gp62 and gp63 hits were manually inspected to find missing protein pairs and accessions or coordinates were added to Table S3. All gp62 hits were inspected for an extended N-terminal sequence missed in many annotations. The resulting sequences were aligned using CLC Main Workbench 25.0.1 Manual adjustments were made to an initial alignment performed with gap open cost 10, gap extension cost 1, and end gap cost free (very accurate alignment setting) settings.

Structure prediction used the AlphaFold3 Server (https://alphafoldserver.com/) and ChimeraX 1.9 (64, 65). Comprehensive topological and domain assignment was performed by integrating multiple prediction algorithms. Transmembrane helices were identified using TMHMM 2.0 while N-terminal signal peptides and their associated cleavage sites were predicted with SignalP 6.0 (66, 67). SignalP was run selecting “other” for organism on the slow model mode. Unless otherwise stated, all bioinformatic analyses used default parameters.

### Figure preparation

Data were graphed using Microsoft Excel or a custom R Shiny application developed with assistance from artificial intelligence (Claude Sonnet 3.7 by Anthropic & Grok 3 by xAI). The Shiny application was built using R v4.5.0 with the following packages: shiny, tidyverse, ggpubr, gridExtra, scales, ggrepel, ggprism, svglite, and jsonlite. The code and instructions for its use are available at (https://github.com/mbaffour/N4-Lysis-paper-codes). Graphics for the figures were assembled in Inkscape 1.3.2 (https://inkscape.org/).

## Acknowledgements

This work was supported by startup funds to J.R. from the Biology Department and the College of Arts & Sciences at Texas A&M University. Research reported in this publication was also supported by the National Institute of General Medical Sciences of the National Institutes of Health under award number R35GM155289 to J.R.. We extend special thanks to Ryland Young for initial support starting this project under Public Health Service grant GM136396 and the Center for Phage Technology, which is jointly supported by Texas A&M AgriLife Research and Texas A&M University. Technical assistance from Gillian Brown and Shivani Bhut as well as productive discussions with all members of the Ramsey lab at Texas A&M and former Young lab members were helpful in the project. We acknowledge Ian Molineux for providing the phage N4 stocks.

Molecular graphics and analyses were performed with UCSF ChimeraX, developed by the Resource for Biocomputing, Visualization, and Informatics at the University of California, San Francisco, with support from National Institutes of Health R01-GM129325 and the Office of Cyber Infrastructure and Computational Biology, National Institute of Allergy and Infectious Diseases. The authors acknowledge the support of the Freiburg Galaxy Team and Björn Grüning, Bioinformatics, University of Freiburg (Germany), funded by the German Federal Ministry of Education and Research BMFTR grant 031 A538A de.NBI-RBC and the Ministry of Science, Research and the Arts Baden-Württemberg (MWK) within the framework of LIBIS/de.NBI Freiburg for usegalaxy.eu resources.

M.B.A., C.C.M, and J.R.R. conceptualized the studies, curated, analyzed, and visualized the data, and supervised all experiments; C.C.M. developed methodology; M.B.A and C.C.M. generated software for data visualization; J.H.S. performed experimental validation; M.B.A and J.R.R. wrote the original manuscript draft; all authors participated in the experimental investigation and reviewing and finalizing the manuscript.

---

**Figure S1.**
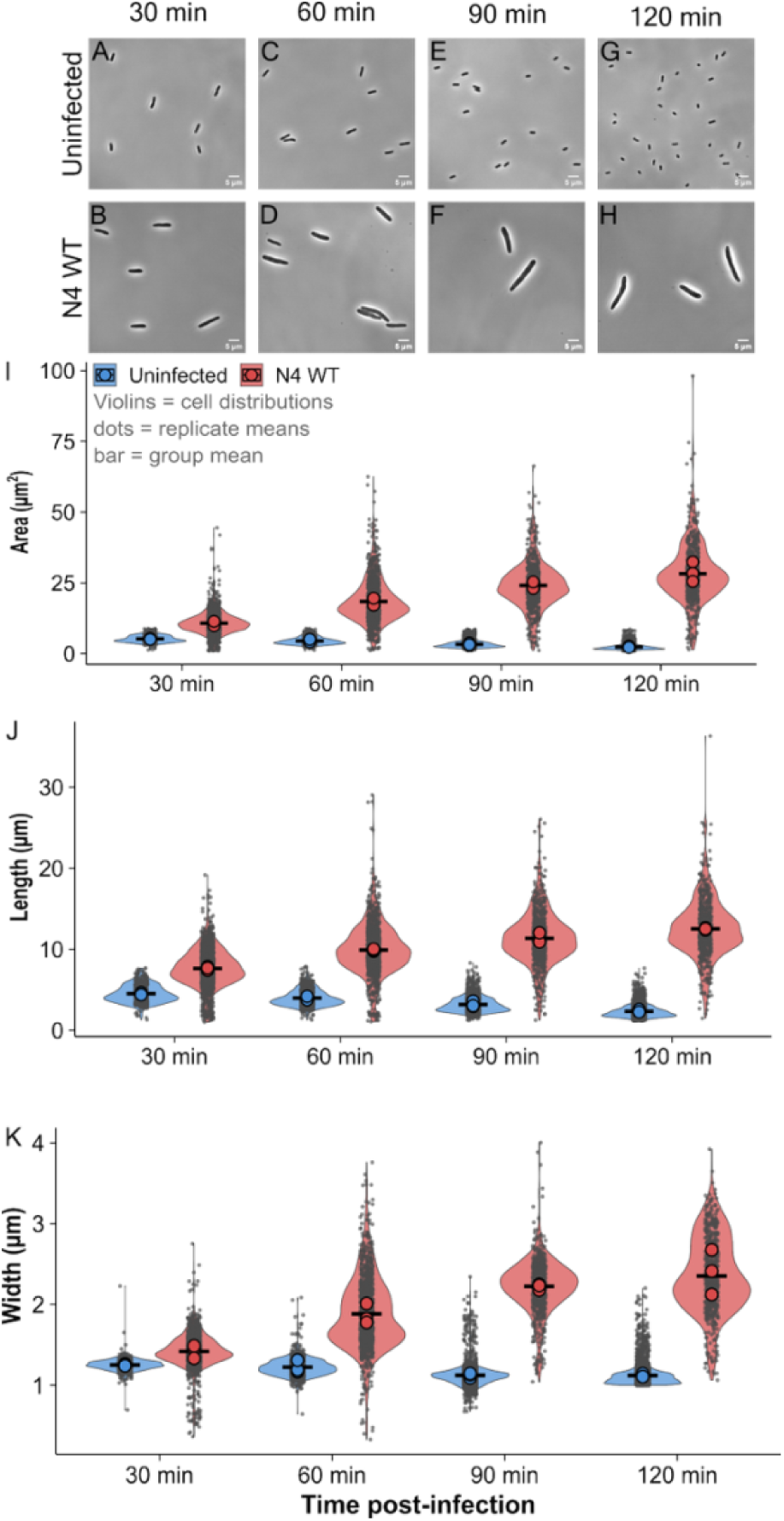
N4-infected cells are significantly larger than uninfected cells. A-H) Representative phase-contrast micrographs of uninfected *E. coli* MG1655 and MOI=5 N4 WT-infected cells at 30 (n=571, 1541), 60 (n=1030, 1075), 90 (n=2743, 715), and 120 (n=3629, 601) min post-infection. Scale bars, 5 µm. I-K) Violin plots showing single-cell distributions of area (µm²), length (µm), and width (µm) for uninfected (blue) and N4 WT-infected (red) cells. Violins represent the full distribution of individual cell measurements; small grey dots indicate individual cells measurements; large colored dots indicate replicate medians; black cross markers indicate the group mean. All measurements are reported as mean ± SD for n = 3 independent biological replicates.

**Figure S2.**
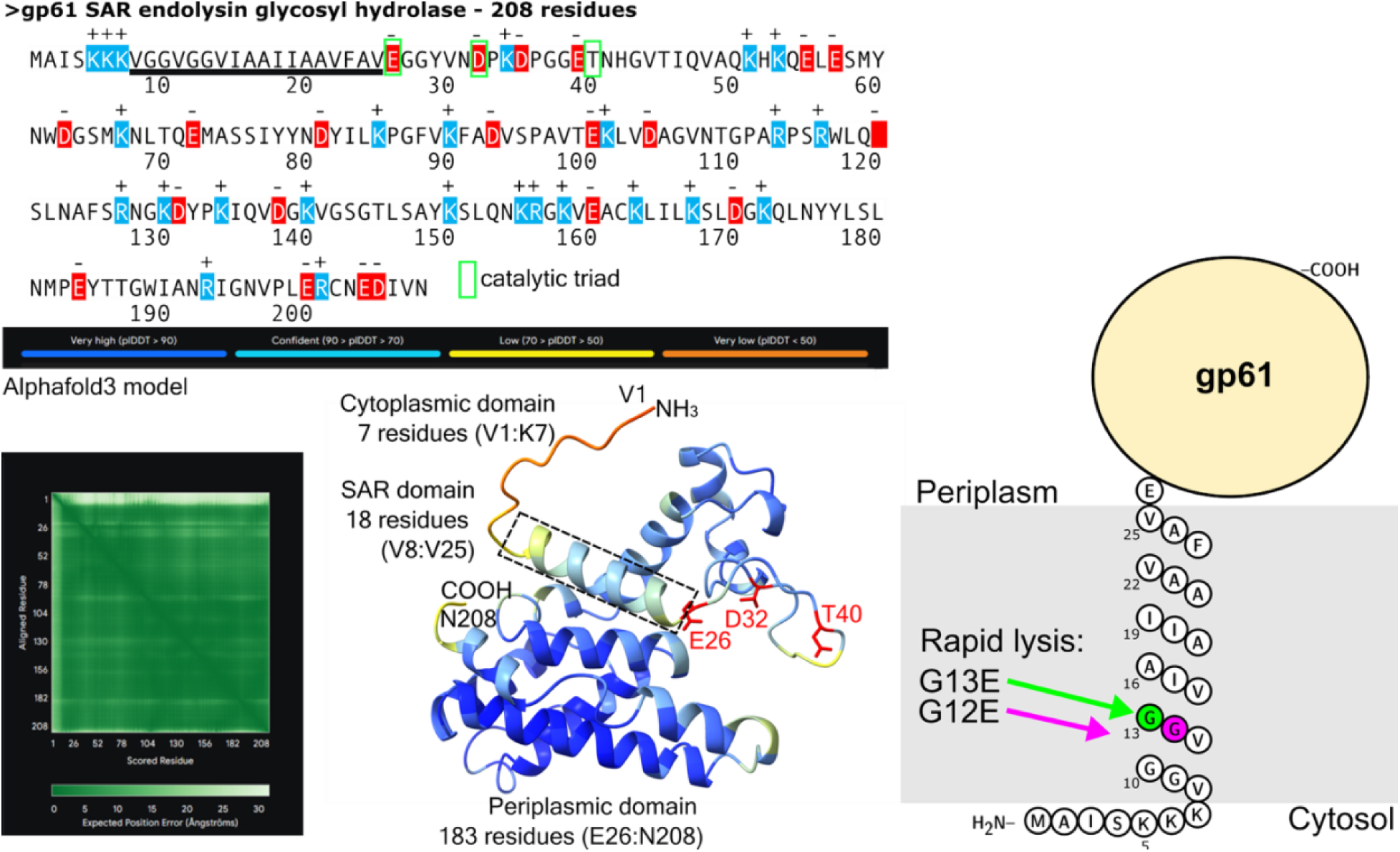
Predicted three-dimensional structure and topological organization of N4 SAR endolysin. The full primary sequence of gp61 with charged residues marked, catalytic triad residues boxed in neon green, and the SAR domain underlined. A tertiary fold model was predicted using the AlphaFold3 Server (top model shown), with the SAR domain boxed, catalytic residues in red, and subfeature lengths annotated. The full plDDT score color scheme displayed on the top model is shown with its expected position error plot. For this model, the ipTM = -pTM = 0.86. At right, a snake plot shows the two positions within the gp61 SAR domain where negatively-charged glutamate residues replaced glycine, leading to rapid lysis.

**Figure S3.**
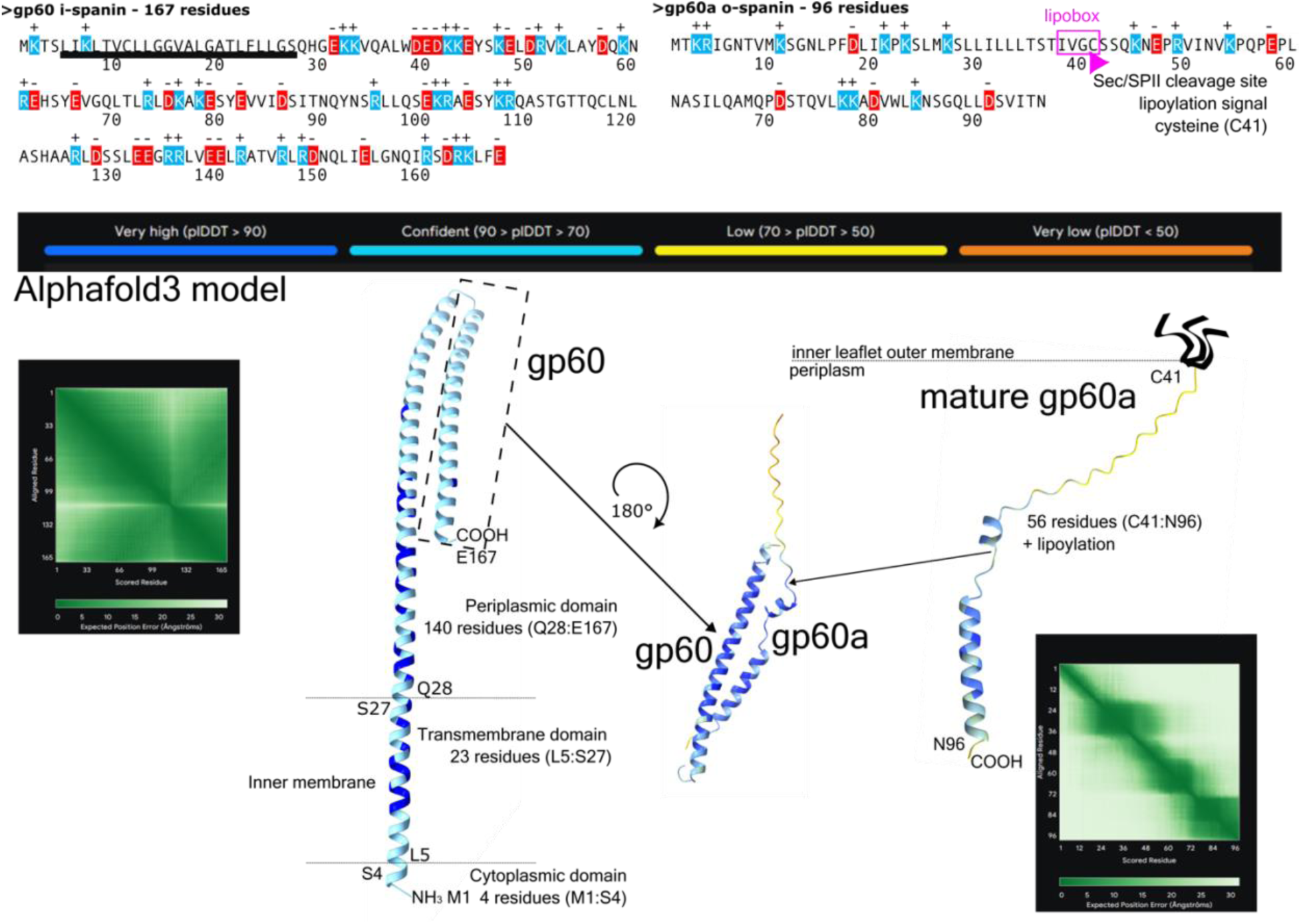
Predicted three-dimensional structures and topological organization of N4 spanins. The full primary sequences of gp60 and gp60a with charged residues marked, lipobox boxed, and the transmembrane domain underlined. Models for each mature protein and a dimer were predicted using the AlphaFold3 Server (top model shown), oriented relative to the respective membrane, and annotated with lengths of subfeatures, and three acylations represented for gp60a. Membrane topology was derived from TMHMM 2.0 (transmembrane helices) and SignalP 6.0 (signal peptides and cleavage sites). The full plDDT score color scheme displayed on the top models is shown with their expected position error plots. For these models, the ipTM = -pTM = 0.61 and ipTM = -pTM = 0.24 for gp60 and gp60a, respectively.

**Figure S4.**
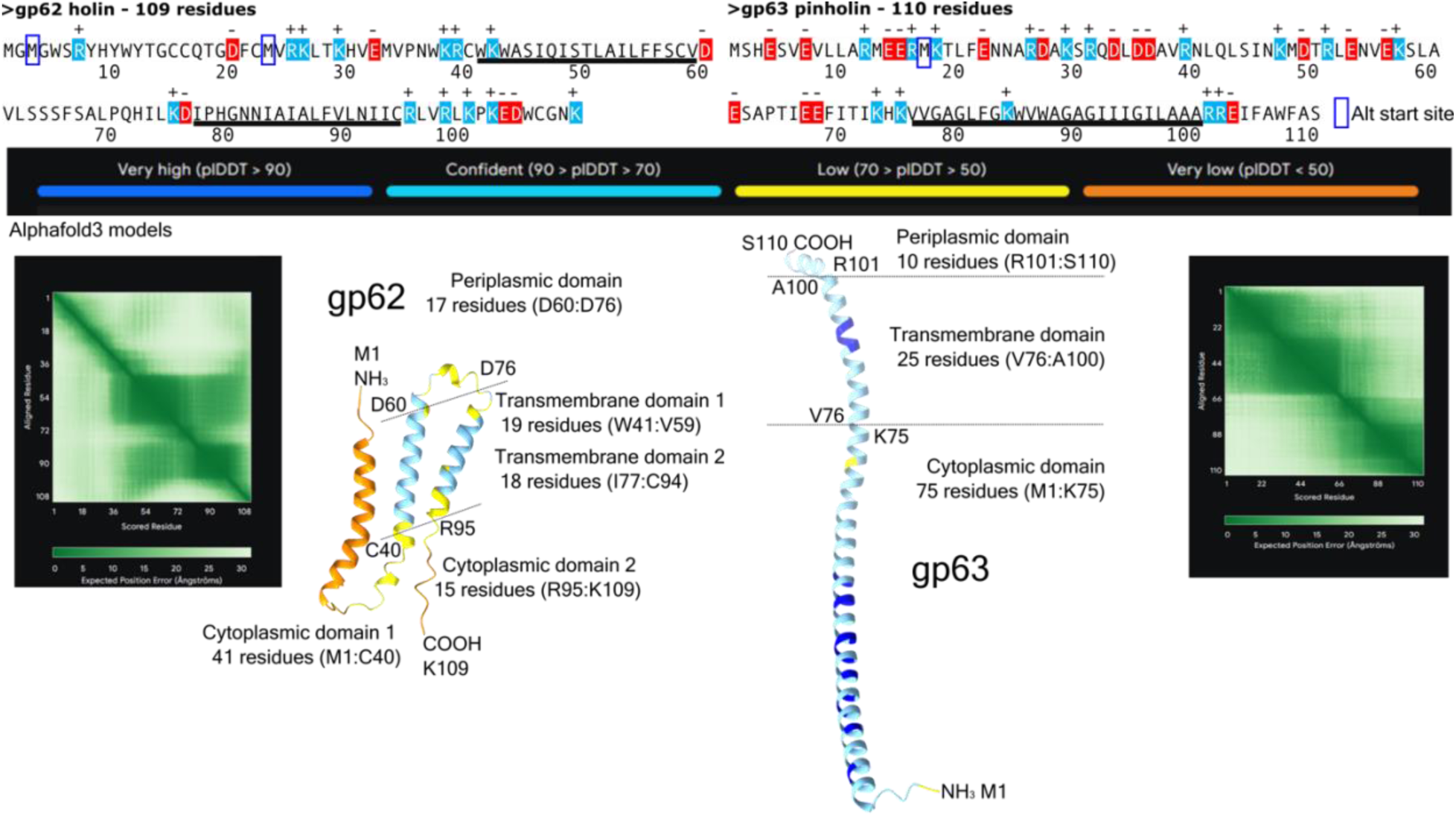
Predicted three-dimensional structures and topological organization of N4 pinholin and holin proteins. The full primary sequences of gp62 and gp63 with charged residues highlighted, predicted transmembrane domains underlined, and alternative start sites boxed. Models for each protein were predicted using AlphaFold3 Server, oriented relative to the respective membrane, and annotated with subfeature lengths. Membrane topology was derived from TMHMM 2.0 (transmembrane helices). The full plDDT score color scheme displayed on the top models is shown with their expected position error plots. For these models, the ipTM = -pTM = 0.35 and ipTM = -pTM = 0.41 for gp62 and gp63, respectively.

**Figure S5.**
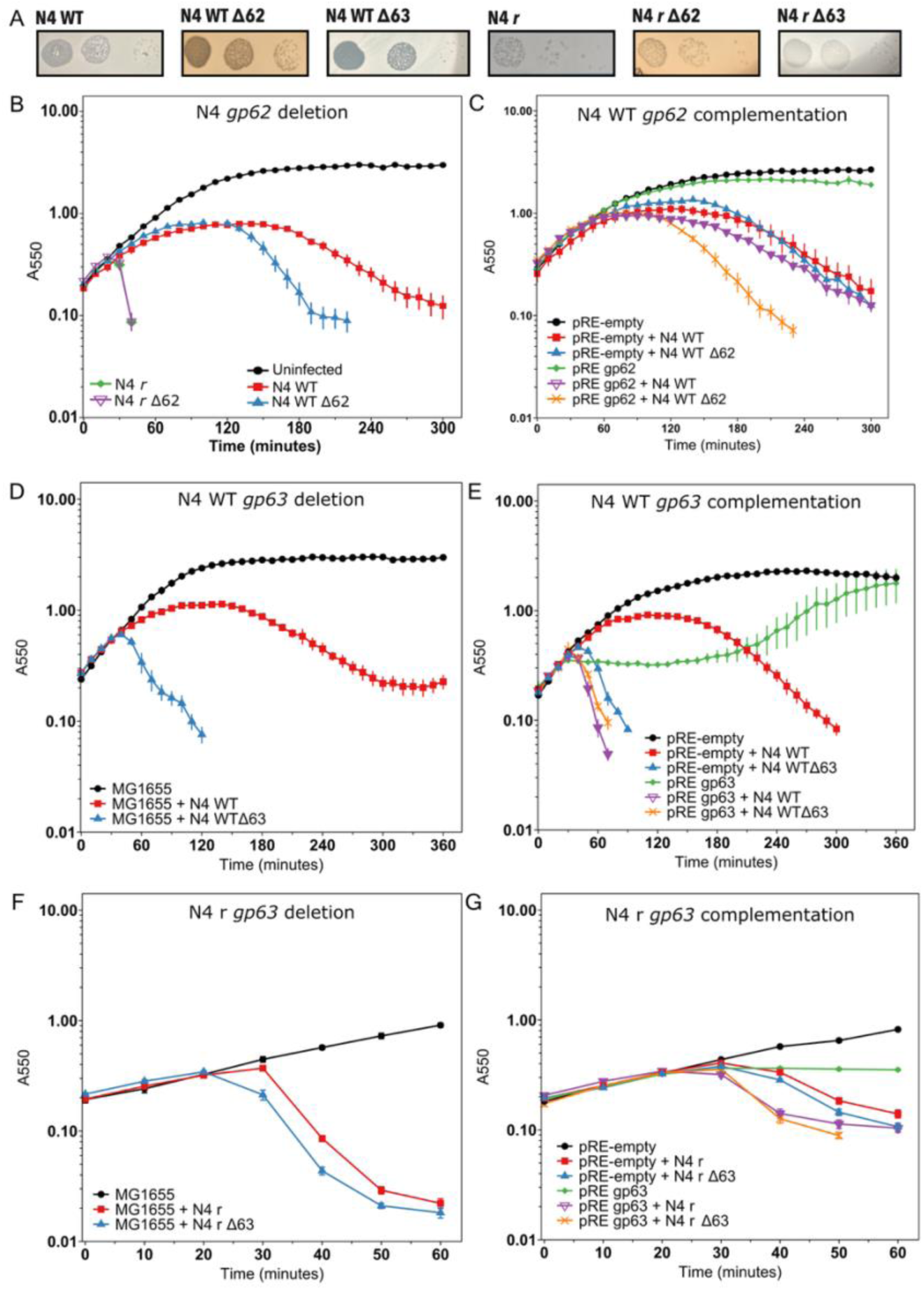
Deletion of gp62 and gp63 in the N4 phage. **A)** Dilution series of N4 plaques for the WT parent and *r* rapid lyser parent backgrounds and their Δ62 and Δ63 derivatives. **B)** MG1655 (solid black circles) infection with N4 at an MOI=5 was monitored for WT (solid red square), the *r* rapid lyser (open purple inverted triangle), and their Δ62 derivatives (solid blue triangle and green diamond, respectively). **C)** Similar to B, with added pQ-Kan and pRE-empty or pRE gp62 induced at 1 mM IPTG. **D)** MG1655 (solid black circles) infection with N4 at an MOI=5 was monitored for WT (solid red square) and its Δ63 derivative (solid blue triangle, from independent replicate number 2 of Table S2). **E)** Same as in D, with added pQ-Kan and pRE-empty or pRE gp63 induced at 1 mM IPTG. **F)** MG1655 (solid black circles) infection with N4 at an MOI=5 was monitored for *r* (solid red square) and its Δ63 derivative (solid blue triangle, from independent replicate number 7 of Table S2). **G)** Same as in E, except infections were with N4 *r* and its derivative Δ63. Lysis curves represent at least three biological replicates and data are presented as the mean ± standard error of the mean.

**Figure S6.**
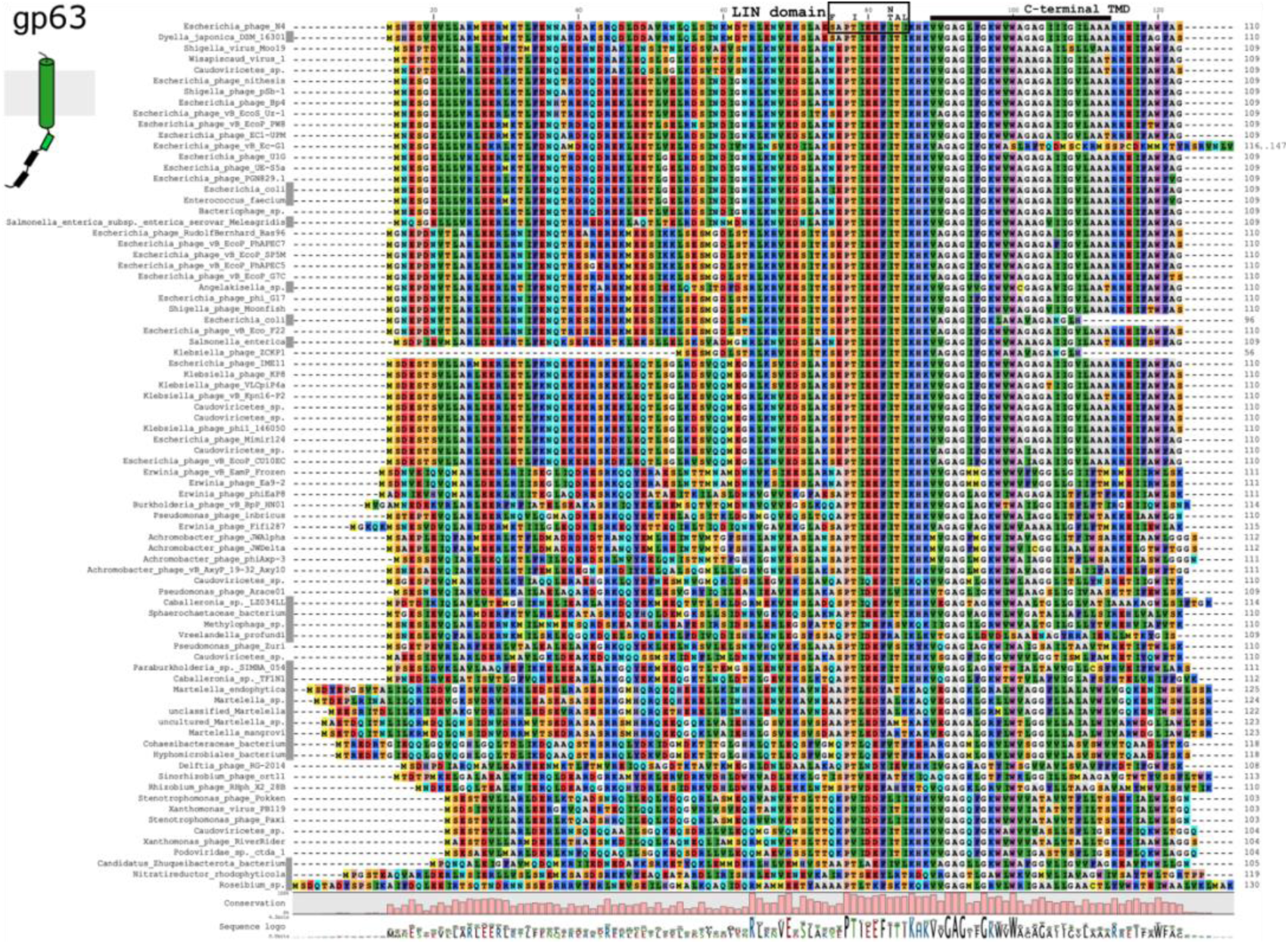
Gp63 pinholin protein alignment. Multiple sequence alignment of 80 proteins with significant sequence similarity to the N4 pinholin gp63 by BLASTp and BLASTx. Sequences in Rasmol colors are labeled by their source organism with bacteria marked by a gray box after the organism name. Predicted coiled-coil regions and the C-terminal anchor transmembrane domain (TMD) are delineated. The LIN domain outlines the cluster of alleles found associated with rapid lysis.

**Figure S7.**
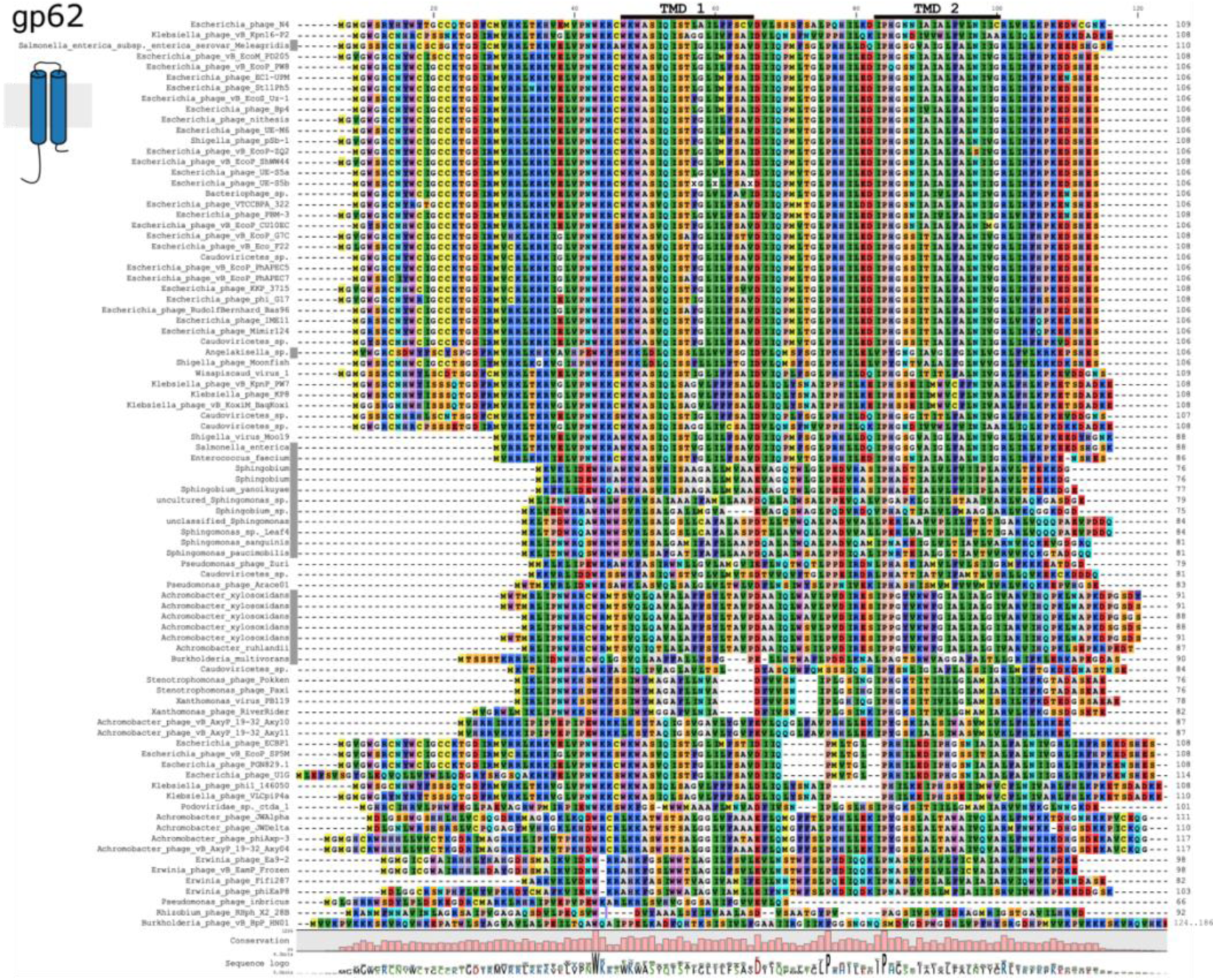
Gp62 holin protein alignment. Multiple sequence alignment of 86 proteins with significant sequence similarity to the N4 gp62 holin by BLASTp and BLASTx. Sequences in Rasmol colors are labeled by their source organism with bacteria marked by a gray box after the organism name. Predicted transmembrane domains (TMD) are delineated.

**Movies S1 and S2. N4 wildtype infection lysis videos**. MG1655 was infected with N4 WT at an MOI=5 and aerated at 37°C. Infected cells were imaged at 100x magnification in time series ∼180 minutes post-infection. Videos are rendered at 11 fps.

**Table S1.**
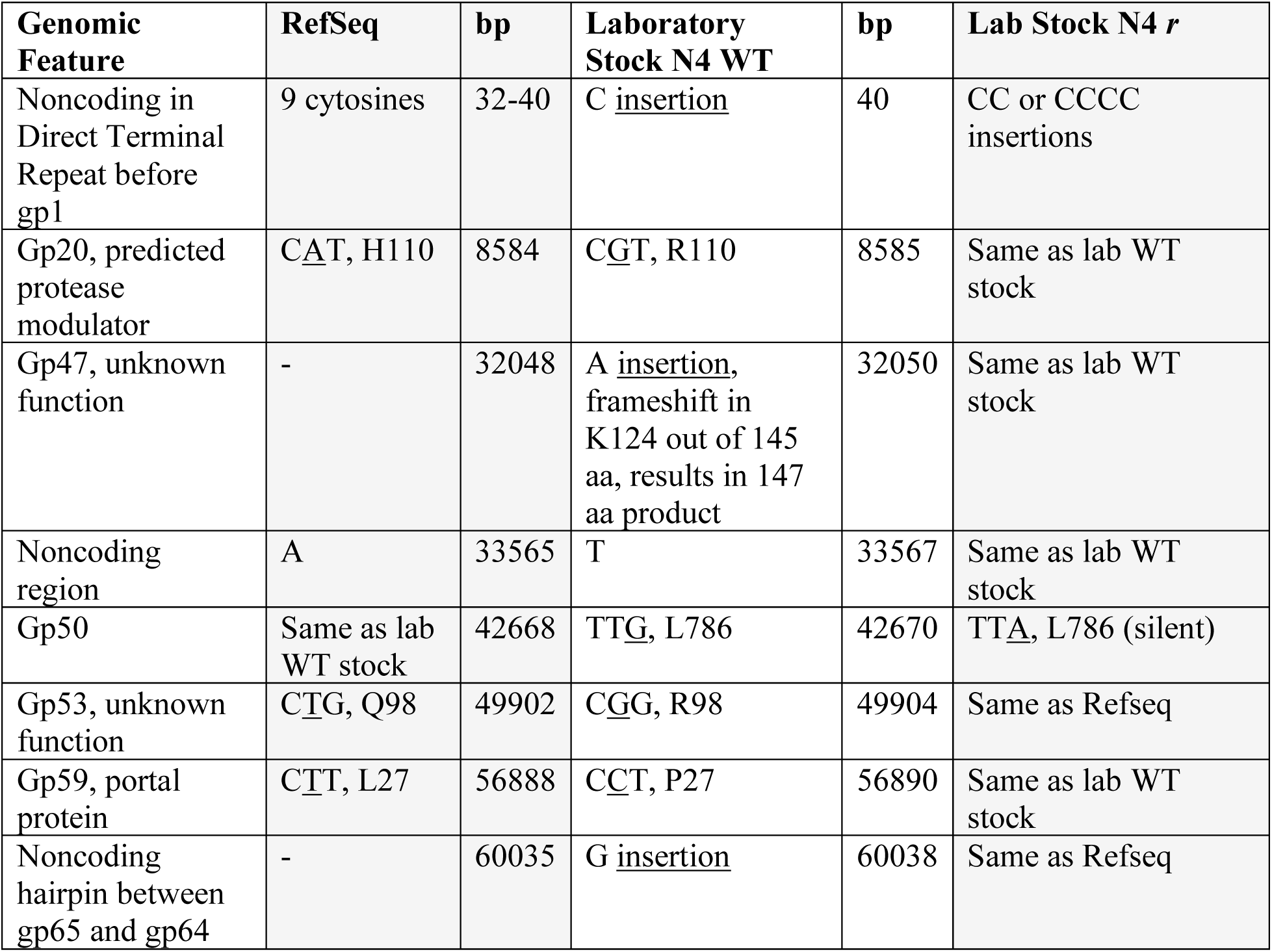
Table describing nucleotide and coding differences between Refseq, lab stock N4 WT, and lab stock N4 *r* phage genome sequences. The lab N4 stocks (WT, 70,156 bp and two *r* derivatives) sequenced via Illumina technology were compared to the NCBI Refseq sequence (NC_008720.1, 70,153 bp) derived from (Genbank EF056009).

**Table S2.**
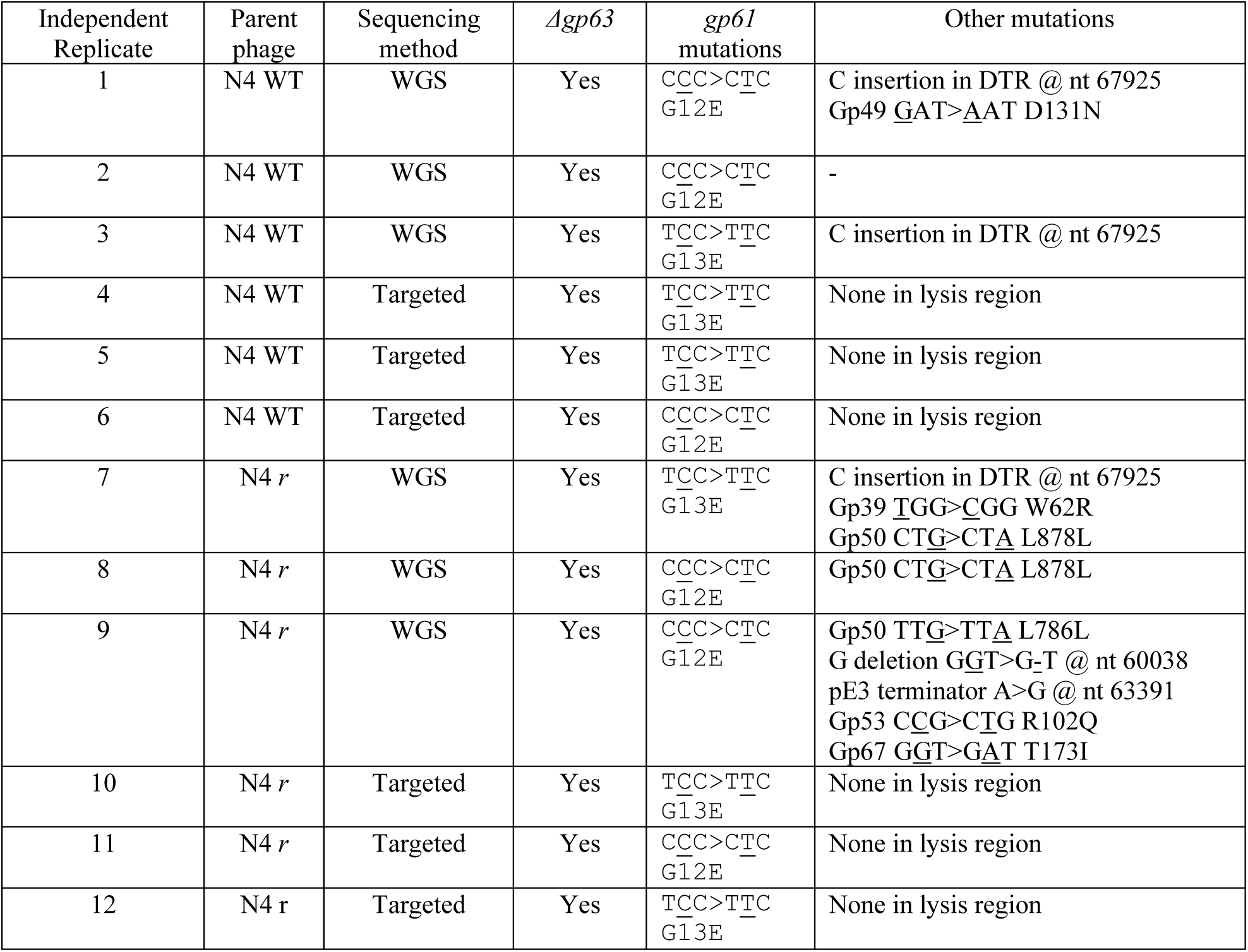
Summary of the sequencing performed on recombinant N4 gp63 deletion strains. Whole genome sequencing (WGS) or targeted sequencing of the lysis gene region were used to assess the deletion of *gp63* from the N4 genome. For the targeted sequencing, the entire lysis region was amplified and sequenced. For the WGS samples, changes found outside the lysis gene region are also listed. A -means no other changes were detected.

| Independent Replicate | Parent phage | Sequencing method | *Δgp63* | *gp61* mutations | Other mutations |
| --- | --- | --- | --- | --- | --- |
| 1 | N4 WT | WGS | Yes | CCC>CTC<br>G12E | C insertion in DTR @ nt 67925<br>Gp49 GAT>AAT D131N |
| 2 | N4 WT | WGS | Yes | CCC>CTC<br>G12E | - |
| 3 | N4 WT | WGS | Yes | TCC>TTC<br>G13E | C insertion in DTR @ nt 67925 |
| 4 | N4 WT | Targeted | Yes | TCC>TTC<br>G13E | None in lysis region |
| 5 | N4 WT | Targeted | Yes | TCC>TTC<br>G13E | None in lysis region |
| 6 | N4 WT | Targeted | Yes | CCC>CTC<br>G12E | None in lysis region |
| 7 | N4 *r* | WGS | Yes | TCC>TTC<br>G13E | C insertion in DTR @ nt 67925<br>Gp39 TGG>CGG W62R<br>Gp50 CTG>CTA L878L |
| 8 | N4 *r* | WGS | Yes | CCC>CTC<br>G12E | Gp50 CTG>CTA L878L |
| 9 | N4 *r* | WGS | Yes | CCC>CTC<br>G12E | Gp50 TTG>TTA L786L<br>G deletion GGT>G-T @ nt 60038<br>pE3 terminator A>G @ nt 63391<br>Gp53 CCG>CTG R102Q<br>Gp67 GGT>GAT T173I |
| 10 | N4 *r* | Targeted | Yes | TCC>TTC<br>G13E | None in lysis region |
| 11 | N4 *r* | Targeted | Yes | CCC>CTC<br>G12E | None in lysis region |
| 12 | N4 *r* | Targeted | Yes | TCC>TTC<br>G13E | None in lysis region |

**Table S3.**
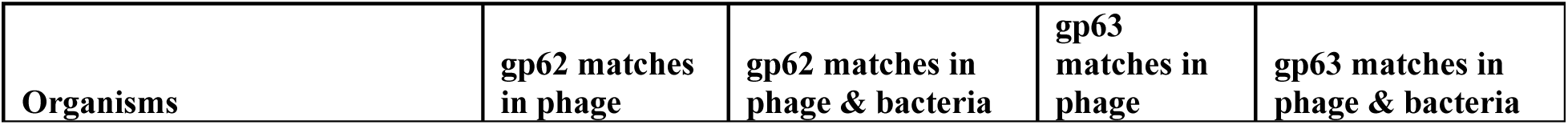
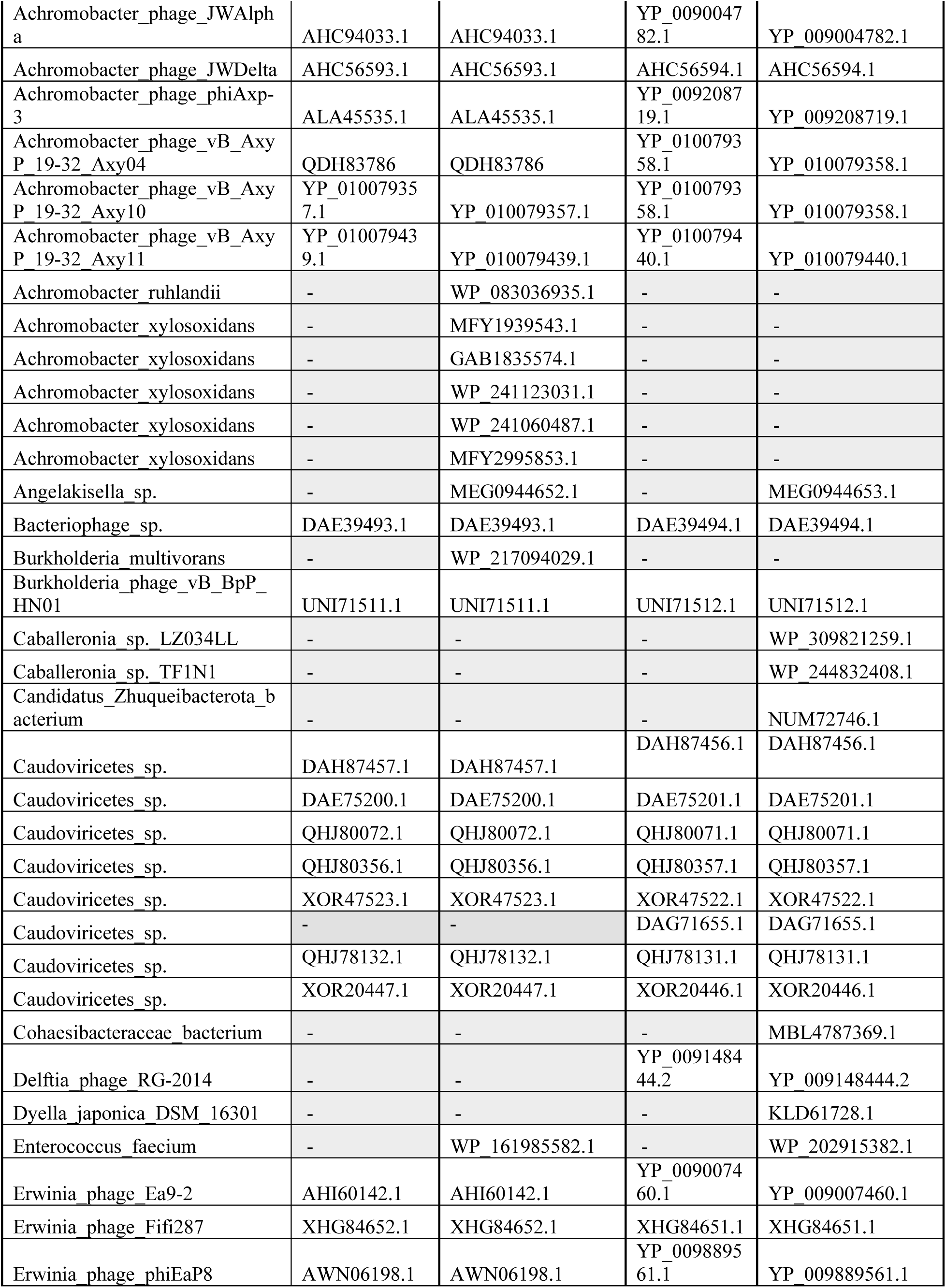
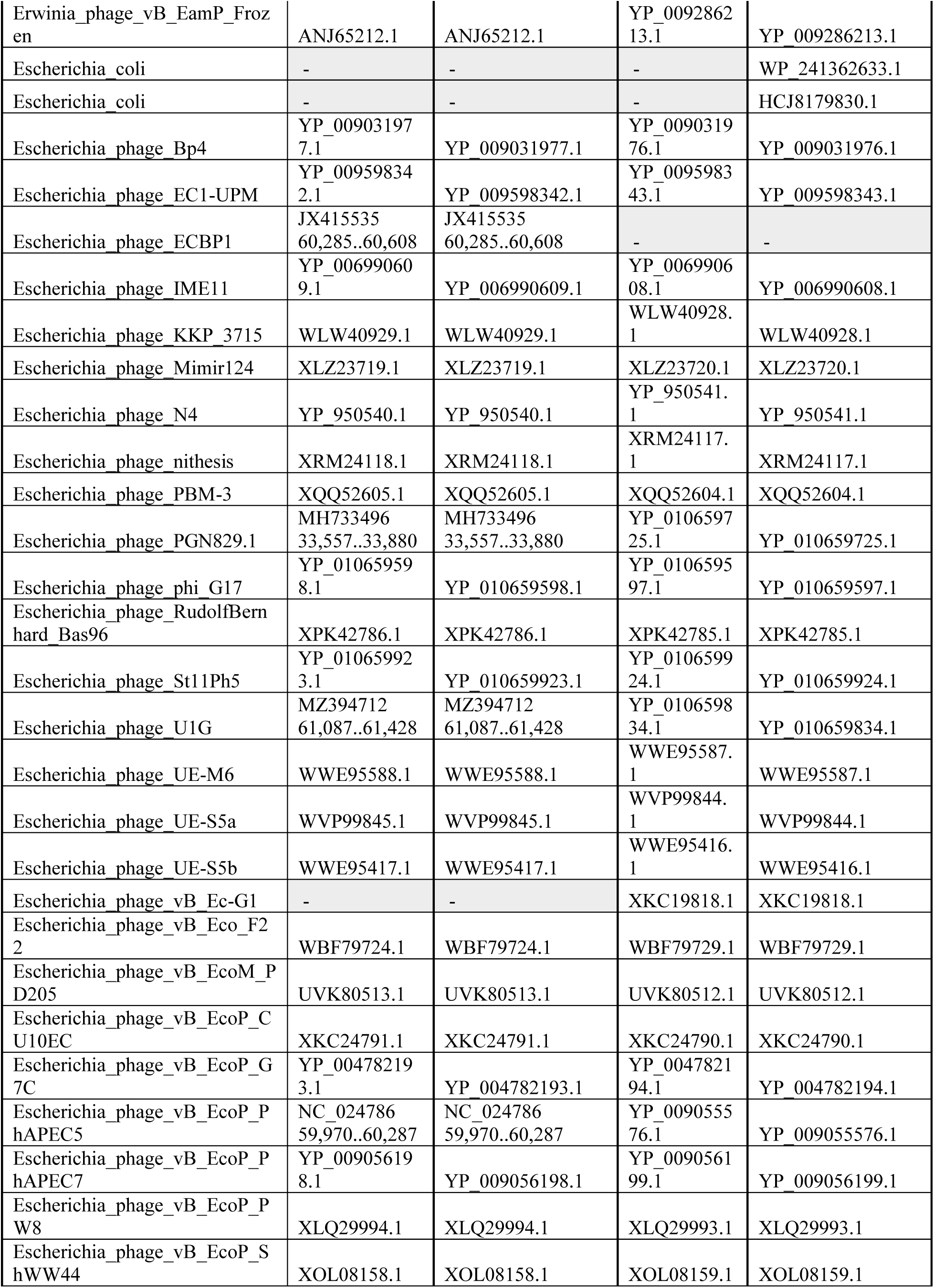
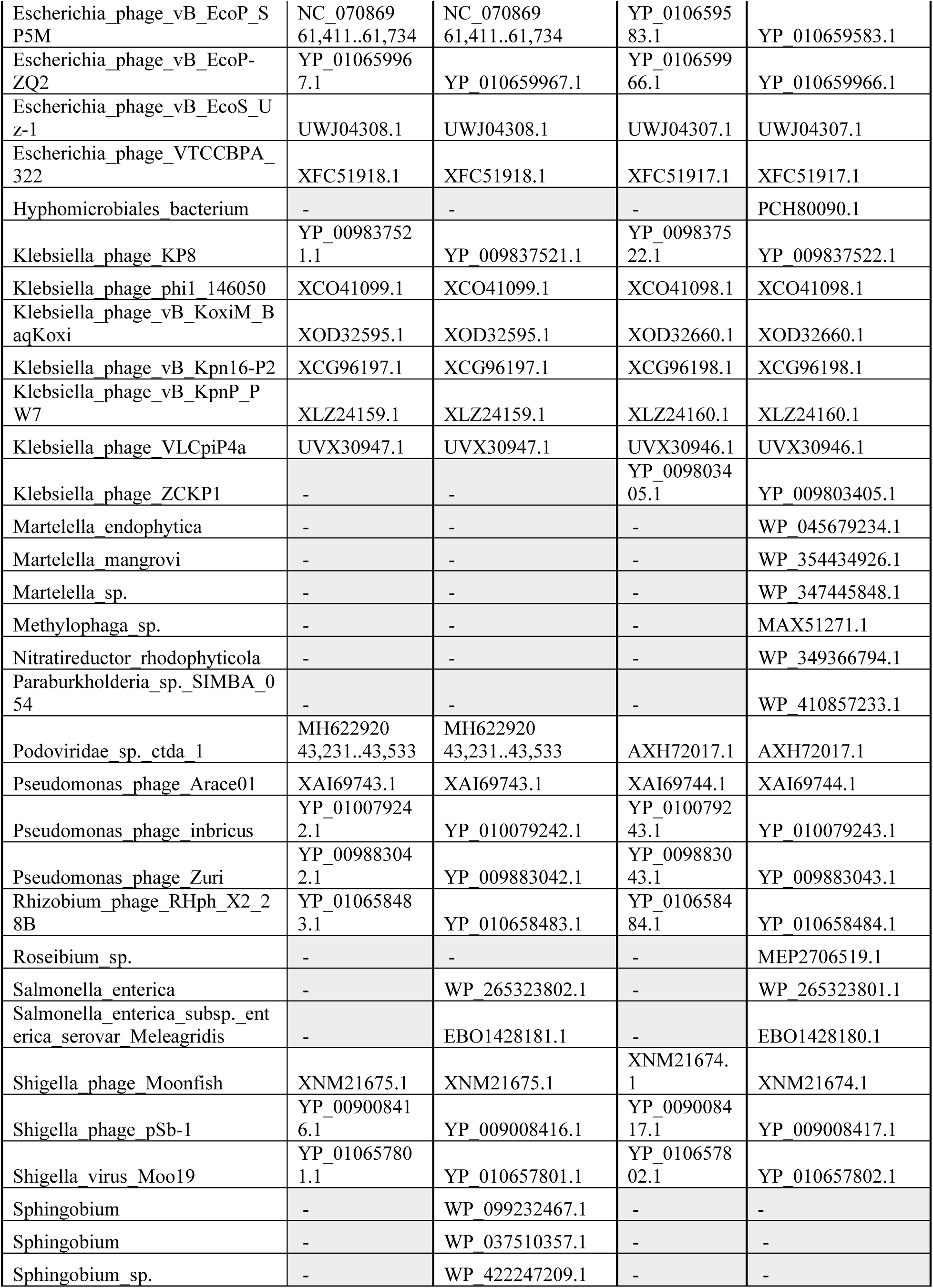
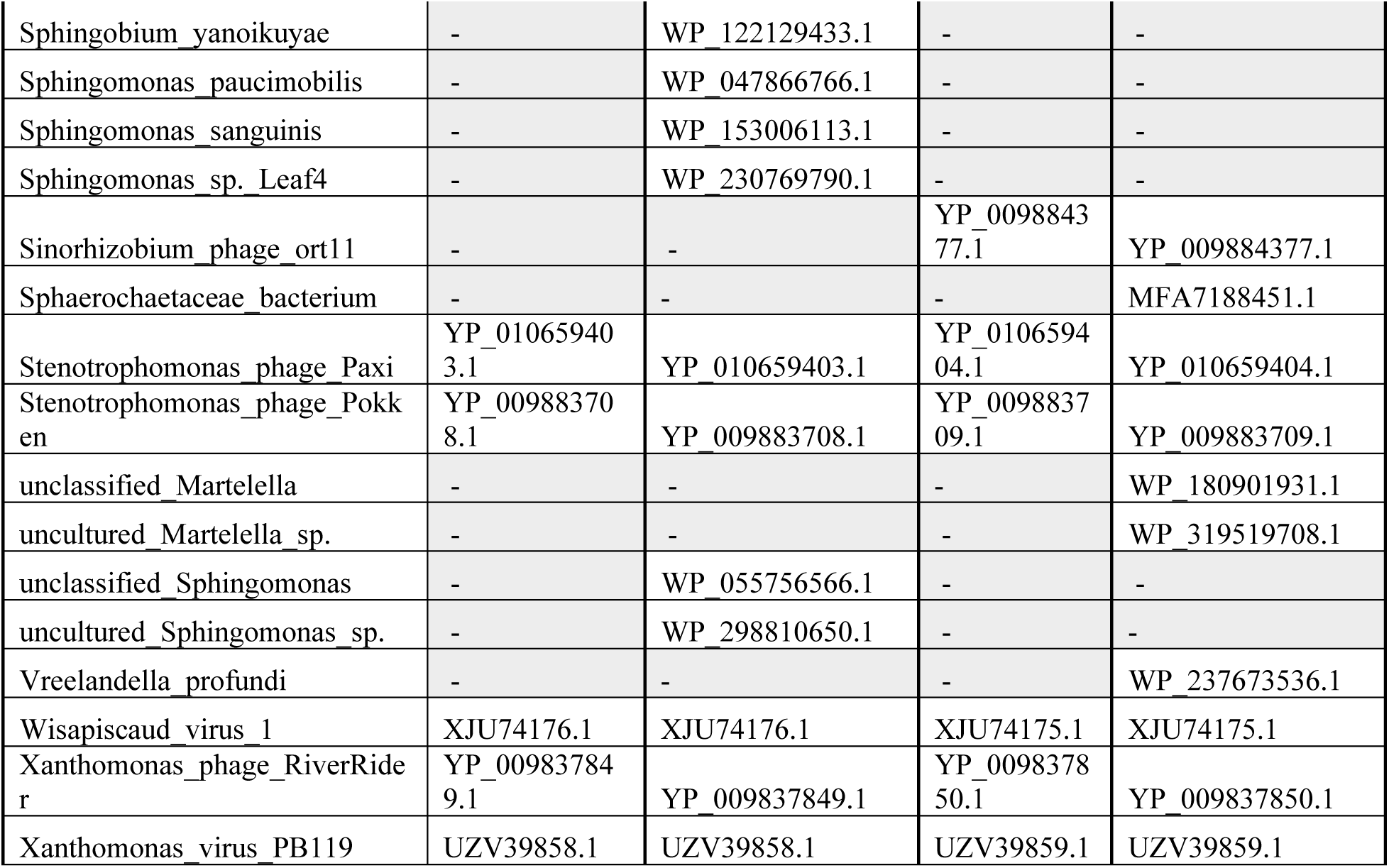
Accessions of gp62 and gp63 proteins hits. Genbank protein accessions for BLASTp and BLASTx hits when searched against a phage only or bacterial and phage database subset and manually added proteins. For unannotated genes, the genome accession is given with the coordinates of the aligned ORF. Due to deduplication to remove 100% identical sequences, not all proteins in the table are shown in alignments.

## References

1. Cahill J, Young R. 2021. Encyclopedia of Virology, p. 501–518. In DH, B, M, Z (eds.), Release of Phages From Prokaryotic Cells, 4th ed. Oxford: Academic Press.

2. Kannoly S, Singh A, Dennehy JJ. 2022. An optimal lysis time maximizes bacteriophage fitness in quasi-continuous culture. Mbio 13:e0359321.

3. Young R. 2013. Phage lysis: do we have the hole story yet? Curr Opin Microbiol 16:790–797.

4. Wang I-N. 2006. Lysis timing and bacteriophage fitness. Genetics 172:17–26.

5. Smith A, Hunter M, Bakshi S, Fusco D. 2026. Live and let die: Lysis time variability and resource limitation shape lytic bacteriophage fitness. PLOS Comput Biol 22:e1013340.

6. Lenneman BR, Rothman-Denes LB. 2015. Structural and biochemical investigation of bacteriophage N4-encoded RNA polymerases. Biomol 5:647–667.

7. Schito GC, Molina AM, Pesce A. 1967. Lysis and Lysis Inhibition with N4 Coliphage. Giornale di microbiologia 15:229–244.

8. Bellis NF, Lokareddy RK, Pavlenok M, Jacobs RQ, Horton SLC, Kizziah JL, Forti F, Schneider DA, Niederweis M, Briani F, Cingolani G. 2026. Structural choreography of bacteriophage N4 ejection proteins and the giant virion-associated RNA polymerase. Nat Commun 10.1038/s41467-026-74701-w.

9. Cahill J, Young R. 2019. Advances in Virus Research, p. 33–70. In Ch 2 Phage lysis: multiple genes for multiple barriers. Academic Press, Elsevier.

10. Kongari R, Rajaure M, Cahill J, Rasche E, Mijalis E, Berry J, Young R. 2018. Phage spanins: diversity, topological dynamics and gene convergence. BMC Bioinformatics 19:326.

11. Catalão MJ, Pimentel M. 2018. Mycobacteriophage Lysis Enzymes: Targeting the Mycobacterial Cell Envelope. Viruses 10:428.

12. Holt A, Cahill J, Ramsey J, Martin C, O’Leary C, Moreland R, Maddox LT, Galbadage T, Sharan R, Sule P, Cirillo JD, Young R. 2021. Phage-Encoded Cationic Antimicrobial Peptide Required for Lysis. J Bacteriol 204:e00214–21.

13. McKitterick AC, Lyerly EW, Bernhardt TG. 2026. Bacteriophages target membrane-anchored glycopolymers to promote host cell lysis and progeny release. Proc Natl Acad Sci United States Am 123:e2606687123.

14. Savva CG, Dewey JS, Moussa SH, To KH, Holzenburg A, Young R. 2014. Stable micron-scale holes are a general feature of canonical holins. Molecular Microbiology 91:57–65.

15. Dewey JS, Savva CG, White RL, Vitha S, Holzenburg A, Young R. 2010. Micron-scale holes terminate the phage infection cycle. Proc National Acad Sci USA 107:2219–2223.

16. Rajaure M, Berry J, Kongari R, Cahill J, Young R. 2015. Membrane fusion during phage lysis. Proc National Acad Sci 112:5497–5502.

17. Pang T, Savva CG, Fleming KG, Struck DK, Young R. 2009. Structure of the lethal phage pinhole. Proc National Acad Sci 106:18966–18971.

18. Xu M, Struck DK, Deaton J, Wang I-N, Young R. 2004. A signal-arrest-release sequence mediates export and control of the phage P1 endolysin. Proc Natl Acad Sci USA 101:6415–6420.

19. Doermann AH. 1948. Lysis and Lysis Inhibition with Escherichia coli Bacteriophage. J Bacteriol 55:257–276.

20. Ramanculov E, Young R. 2001. Genetic analysis of the T4 holin: timing and topology. Gene 265:25–36.

21. Tran TAT, Struck DK, Young R. 2005. Periplasmic domains define holin-antiholin interactions in T4 lysis inhibition. J Bacteriol 187:6631–6640.

22. Schwarzkopf JMF, Mehner-Breitfeld D, Brüser T. 2024. A dimeric holin/antiholin complex controls lysis by phage T4. Front Microbiol 15:1419106.

23. Burch LH, Zhang L, Chao FG, Xu H, Drake JW. 2011. The Bacteriophage T4 Rapid-Lysis Genes and Their Mutational Proclivities. J Bacteriol 193:3537–3545.

24. Anderson CW, Eigner J. 1971. Breakdown and Exclusion of Superinfecting T-Even Bacteriophage in Escherichia coli. J Virol 8:869–886.

25. Krieger IV, Kuznetsov V, Chang J-Y, Zhang J, Moussa SH, Young RF, Sacchettini JC. 2020. The Structural Basis of T4 Phage Lysis Control: DNA as the Signal for Lysis Inhibition. J Mol Biol 432:4623–4636.

26. Hays SG, Seed KD. 2020. Dominant Vibrio cholerae phage exhibits lysis inhibition sensitive to disruption by a defensive phage satellite. Elife 9:e53200.

27. Stojković EA, Rothman-Denes LB. 2007. Coliphage N4 N-Acetylmuramidase Defines a New Family of Murein Hydrolases. J Mol Biol 366:406–419.

28. Summer EJ, Berry J, Tran TAT, Niu L, Struck DK, Young R. 2007. Rz/Rz1 Lysis Gene Equivalents in Phages of Gram-negative Hosts. J Mol Biol 373:1098–1112.

29. Schito GC. 1967. Intracellular crystallization of the DNA coliphage N4. Virology 32:723– 725.

30. Pesce A, Satta G, Schito GC. 1969. Factors in Lysis-Inhibition by N4 Coliphage. Giornale di microbiologia 17:119–129.

31. Wittmann J, Klumpp J, Switt AIM, Yagubi A, Ackermann H-W, Wiedmann M, Svircev A, Nash JHE, Kropinski AM. 2015. Taxonomic reassessment of N4-like viruses using comparative genomics and proteomics suggests a new subfamily -“Enquartavirinae”. Archives of virology 160:3053–3062.

32. Wittmann J, Turner D, Millard AD, Mahadevan P, Kropinski AM, Adriaenssens EM. 2020. From Orphan Phage to a Proposed New Family–The Diversity of N4-Like Viruses. Antibiotics 9:663.

33. Zheng K, Liang Y, Paez-Espino D, Zou X, Gao C, Shao H, Sung YY, Mok WJ, Wong LL, Zhang Y-Z, Tian J, Chen F, Jiao N, Suttle CA, He J, McMinn A, Wang M. 2023. Identification of hidden N4-like viruses and their interactions with hosts. mSystems e00197–23.

34. Kulikov E, Kropinski AM, Golomidova A, Lingohr E, Govorun V, Serebryakova M, Prokhorov N, Letarova M, Manykin A, Strotskaya A, Letarov A. 2012. Isolation and characterization of a novel indigenous intestinal N4-related coliphage vB_EcoP_G7C. Virology 426:93–99.

35. Wittmann J, Dreiseikelmann B, Rohde M, Meier-Kolthoff JP, Bunk B, Rohde C. 2014. First genome sequences of Achromobacter phages reveal new members of the N4 family. Virology journal 11:14.

36. Menon ND, Kumar MS, Babu TGS, Bose S, Vijayakumar G, Baswe M, Chatterjee M, D’Silva JR, Shetty K, Haripriyan J, Kumar A, Nair S, Somanath P, Nair BG, Nizet V, Kumar GB. 2021. A Novel N4-Like Bacteriophage Isolated from a Wastewater Source in South India with Activity against Several Multidrug-Resistant Clinical Pseudomonas aeruginosa Isolates. Msphere 6:e01215–20.

37. Barenboim M, Chang CY, Hajj F dib, Young R. 1999. Characterization of the dual start motif of a class II holin gene. Mol Microbiol 32:715–727.

38. Grundling A, Smith DL, Bläsi U, Young R. 2000. Dimerization between the holin and holin inhibitor of phage lambda. J Bacteriol 182:6075–6081.

39. Park T, Struck DK, Deaton JF, Young R. 2006. Topological dynamics of holins in programmed bacterial lysis. Proc National Acad Sci 103:19713–19718.

40. Bläsi U, Nam K, Hartz D, Gold L, Young R. 1989. Dual translational initiation sites control function of the lambda S gene. EMBO J 8:3501–3510.

41. Bläsi U, Chang CY, Zagotta MT, Nam KB, Young R. 1990. The lethal lambda S gene encodes its own inhibitor. EMBO J 9:981–989.

42. EA S. 2005. Dissertation: Characterization of the Coliphage N4-encoded N-Acetylmuramidase, a member of a new family of peptidoglycan-hydrolyzing enzymes. ProQuest Information and Learning Company, The University of Chicago.

43. Sun Q, Kuty GF, Arockiasamy A, Xu M, Young R, Sacchettini JC. 2009. Regulation of a muralytic enzyme by dynamic membrane topology. Nat Struct Mol Biol 16:1192–1194.

44. Lu M-J, Henning U. 1994. Superinfection exclusion by T-even-type coliphages. Trends Microbiol 2:137–139.

45. Kanamaru S, Uchida K, Nemoto M, Fraser A, Arisaka F, Leiman PG. 2020. Structure and Function of the T4 Spackle Protein Gp61.3. Viruses 12:1070.

46. Mehner-Breitfeld D, Schwarzkopf JMF, Young R, Kondabagil K, Brüser T. 2021. The Phage T4 Antiholin RI Has a Cleavable Signal Peptide, Not a SAR Domain. Front Microbiol 12:712460.

47. Schwarzkopf JMF, Viveros RP, Burdur AN, Mehner-Breitfeld D, Tschowri N, Brüser T. 2025. The short cytoplasmic region of phage T4 holin is essential for the transition from impermeable membrane protein complexes to permeable pores. Front Microbiol 16:1579756.

48. Manohar P, Oppong K, Young R. 2026. Characterization of Two Salmonella Myoviruses and a Novel Lysis-Inhibition System. PHAGE: Ther, Appl, Res 10.1177/26416549261444851.

49. Gromkova RH. 1968. T-related Bacteriophages Isolated from Shigella sonnei. J Virol 2:692– 694.

50. Schito GC. 1974. Development of coliphage N4: ultrastructural studies. J Virol 13:186–196.

51. Kannoly S, Oken G, Shadan J, Musheyev D, Singh K, Singh A, Dennehy JJ. 2022. Single-Cell Approach Reveals Intercellular Heterogeneity in Phage-Producing Capacities. Microbiol Spectr 11:e02663–21.

52. Adler BA, Hessler T, Cress BF, Lahiri A, Mutalik VK, Barrangou R, Banfield J, Doudna JA. 2022. Broad-spectrum CRISPR-Cas13a enables efficient phage genome editing. Nat Microbiol 7:1967–1979.

53. Trofimova E, Jaschke PR. 2021. Plaque Size Tool: An automated plaque analysis tool for simplifying and standardising bacteriophage plaque morphology measurements. Virology 561:1– 5.

54. Shamash M, Maurice CF. 2021. OnePetri: Accelerating Common Bacteriophage Petri Dish Assays with Computer Vision. PHAGE: Ther, Appl, Res 2:224–231.

55. Schindelin J, Arganda-Carreras I, Frise E, Kaynig V, Longair M, Pietzsch T, Preibisch S, Rueden C, Saalfeld S, Schmid B, Tinevez J-Y, White DJ, Hartenstein V, Eliceiri K, Tomancak P, Cardona A. 2012. Fiji: an open-source platform for biological-image analysis. Nat Methods 9:676–682.

56. Lord SJ, Velle KB, Mullins RD, Fritz-Laylin LK. 2020. SuperPlots: Communicating reproducibility and variability in cell biology. J Cell Biol 219:e202001064.

57. Ducret A, Quardokus EM, Brun YV. 2016. MicrobeJ, a tool for high throughput bacterial cell detection and quantitative analysis. Nat Microbiol 1:16077.

58. Corban JE, Ramsey J. 2021. Characterization and complete genome sequence of Privateer, a highly prolate Proteus mirabilis podophage. Peerj 9:e10645.

59. Ramsey J, Rasche H, Maughmer C, Criscione A, Mijalis E, Liu M, Hu JC, Young R, Gill JJ. 2020. Galaxy and Apollo as a biologist-friendly interface for high-quality cooperative phage genome annotation. PLoS Comput Biol 16:e1008214.

60. Community TG, Abueg LAL, Afgan E, Allart O, Awan AH, Bacon WA, Baker D, Bassetti M, Batut B, Bernt M, Blankenberg D, Bombarely A, Bretaudeau A, Bromhead CJ, Burke ML, Capon PK, Čech M, Chavero-Díez M, Chilton JM, Collins TJ, Coppens F, Coraor N, Cuccuru G, Cumbo F, Davis J, Geest PFD, Koning W de, Demko M, DeSanto A, Begines JMD, Doyle MA, Droesbeke B, Erxleben-Eggenhofer A, Föll MC, Formenti G, Fouilloux A, Gangazhe R, Genthon T, Goecks J, Beltran ANG, Goonasekera NA, Goué N, Griffin TJ, Grüning BA, Guerler A, Gundersen S, Gustafsson OJR, Hall C, Harrop TW, Hecht H, Heidari A, Heisner T, Heyl F, Hiltemann S, Hotz H-R, Hyde CJ, Jagtap PD, Jakiela J, Johnson JE, Joshi J, Jossé M, Jum’ah K, Kalaš M, Kamieniecka K, Kayikcioglu T, Konkol M, Kostrykin L, Kucher N, Kumar A, Kuntz M, Lariviere D, Lazarus R, Bras YL, Corguillé GL, Lee J, Leo S, Liborio L, Libouban R, Tabernero DL, Lopez-Delisle L, Los LS, Mahmoud A, Makunin I, Marin P, Mehta S, Mok W, Moreno PA, Morier-Genoud F, Mosher S, Müller T, Nasr E, Nekrutenko A, Nelson TM, Oba AJ, Ostrovsky A, Polunina PV, Poterlowicz K, Price EJ, Price GR, Rasche H, Raubenolt B, Royaux C, Sargent L, Savage MT, Savchenko V, Savchenko D, Schatz MC, Seguineau P, Serrano-Solano B, Soranzo N, Srikakulam SK, Suderman K, Syme AE, Tangaro MA, Tedds JA, Tekman M, Thang WC (Mike), Thanki AS, Uhl M, Beek M van den, Varshney D, Vessio J, Videm P, Kuster GV, Watson GR, Whitaker-Allen N, Winter U, Wolstencroft M, Zambelli F, Zierep P, Zoabi R. 2024. The Galaxy platform for accessible, reproducible, and collaborative data analyses: 2024 update. Nucleic Acids Res 52:W83–W94.

61. Li W, Godzik A. 2006. Cd-hit: a fast program for clustering and comparing large sets of protein or nucleotide sequences. Bioinformatics 22:1658–1659.

62. Fu L, Niu B, Zhu Z, Wu S, Li W. 2012. CD-HIT: accelerated for clustering the next-generation sequencing data. Bioinformatics 28:3150–3152.

63. Camacho C, Coulouris G, Avagyan V, Ma N, Papadopoulos J, Bealer K, Madden TL. 2009. BLAST+: architecture and applications. BMC Bioinform 10:421.

64. Abramson J, Adler J, Dunger J, Evans R, Green T, Pritzel A, Ronneberger O, Willmore L, Ballard AJ, Bambrick J, Bodenstein SW, Evans DA, Hung C-C, O’Neill M, Reiman D, Tunyasuvunakool K, Wu Z, Žemgulytė A, Arvaniti E, Beattie C, Bertolli O, Bridgland A, Cherepanov A, Congreve M, Cowen-Rivers AI, Cowie A, Figurnov M, Fuchs FB, Gladman H, Jain R, Khan YA, Low CMR, Perlin K, Potapenko A, Savy P, Singh S, Stecula A, Thillaisundaram A, Tong C, Yakneen S, Zhong ED, Zielinski M, Žídek A, Bapst V, Kohli P, Jaderberg M, Hassabis D, Jumper JM. 2024. Accurate structure prediction of biomolecular interactions with AlphaFold 3. Nature 630:493–500.

65. Meng EC, Goddard TD, Pettersen EF, Couch GS, Pearson ZJ, Morris JH, Ferrin TE. 2023. UCSF ChimeraX: Tools for structure building and analysis. Protein Sci 32:e4792.

66. Krogh A, Larsson B, Heijne G von, Sonnhammer ELL. 2001. Predicting transmembrane protein topology with a hidden markov model: application to complete genomes11Edited by F. Cohen. J Mol Biol 305:567–580.

67. Teufel F, Armenteros JJA, Johansen AR, Gíslason MH, Pihl SI, Tsirigos KD, Winther O, Brunak S, Heijne G von, Nielsen H. 2022. SignalP 6.0 predicts all five types of signal peptides using protein language models. Nat Biotechnol 40:1023–1025.

68. Zhang N, Young R. 1999. Complementation and characterization of the nested Rz and Rz1 reading frames in the genome of bacteriophage lambda. Mol Gen Genet 262:659–667.

69. AM M, A P, GC S. 1965. Un nuovo batteriofago attivo sul ceppo K12 di E. coli I. Caratteristiche. Bollettino dell’Istituto sieroterapico milanese 329–332.

70. Gründling A, Manson MD, Young R. 2001. Holins kill without warning. Proc National Acad Sci 98:9348–9352.

71. Berry J, Summer EJ, Struck DK, Young R. 2008. The final step in the phage infection cycle: the Rz and Rz1 lysis proteins link the inner and outer membranes. Mol Microbiol 70:341–351.

72. White R, Chiba S, Pang T, Dewey JS, Savva CG, Holzenburg A, Pogliano K, Young R. 2011. Holin triggering in real time. Proc National Acad Sci 108:798–803.

73. Ruegg TL, Pereira JH, Chen JC, DeGiovanni A, Novichkov P, Mutalik VK, Tomaleri GP, Singer SW, Hillson NJ, Simmons BA, Adams PD, Thelen MP. 2018. Jungle Express is a versatile repressor system for tight transcriptional control. Nature communications 9:4596–33.

